# Tolerance of chronic HDACi-administration used for treatment of neurological and visceral disease including lung pathogenesis

**DOI:** 10.1101/191635

**Authors:** Md. Suhail Alam, Bruce Cooper, Kasturi Haldar

## Abstract

Histone deacetylase (HDAC) inhibitors are of significant interest as drugs. However, their use in neurological disorders has raised concern because HDACs are required for brain function. We have previously shown that a triple combination formulation (TCF) of the pan HDACi vorinostat (Vo) improves pharmacokinetic exposure and entry of Vo into the brain. TCF treatment significantly delayed both neurodegeneration and death in the *Npc1*^*nmf164*^ murine model of Niemann Pick Type C (NPC) disease. The TCF induces no metabolic toxicity, but it’s risk to normal brain functions and potential utility in treating lung disease, a major NPC clinical complication, remain unknown. Here we report that TCF administered for 8-10 months was not detrimental to brain or neuromuscular functions of healthy mice, based on quantitative analyses of major Purkinje neurons in the cerebellum, inflammation in the hippocampus and symptoms of progressive neurocognitive/muscular disease. The TCF was also not injurious to lung tissue but rather improved delivery of Vo to lungs and reduced accumulation of foamy macrophages in *Npc1*^*nmf164*^ mice. Together these data support feasibility of tolerable, chronic administration of an HDACi formulation to treat brain and systemic disease including lung pathology, a frequent cause of death in NPC and possibly other neurological diseases.

## Introduction

Histone deacetylase inhibitors (HDACi) are emerging therapeutics for a broad range of diseases including cancer and neurodegeneration ^1-4^. They block HDAC enzymes, to promote acetylation of both histones and non-histone proteins to elicit complex cellular changes ^5,6^. HDACi-induced histone modifications have been shown to increase or decrease transcriptional expression of mutated target gene(s) in many genetic diseases as well as indirect benefit through modulation of chaperone and proteostatic networks ^7-9^. Due to their broad effects on transcription, it is particularly important to maximize HDACi efficacy while limiting dose. We previously reported on development and validation of a therapeutic strategy of a triple combination formulation (TCF) of the HDACi vorinostat (Vo) that enabled lowering concentrations of Vo to treat cerebral disease as well as inflammation in liver and spleen, in a mouse model of a fatal cerebellar disorder Niemann-Pick Type C (NPC) disease ^10^.

NPC is caused by defect in either *Npc1* or *Npc2* genes ^11^. It is a rare autosomal recessive neurodegenerative disease. 95% of cases are due to defect in *Npc1.* Cells with defects in either *Npc* genes accumulate cholesterol late endosomes/lysosomes ^12,13^. A point mutation in *Npc1* gene that blocks cholesterol transport in cells is causative for neurodegeneration in a mouse model ^14^. At the organismal level, in the central nervous system (CNS), *Npc1* is essential for myelination ^15^ and likely additional functions^16^. Neurodegeneration is a hallmark of clinical NPC disease. Disease progression can be heterogeneous and slow but once initiated, is invariably fatal ^11^. Splenomegaly and hepatomegaly are common presenting symptoms in pediatric cases followed by neurocognitive and neuromuscular degeneration ^17^. Lung disease is prominent and can even be cause of death ^18,19^.

Presently the only available treatment for NPC is miglustat (Zavesca^™^), an iminosugar that decreases glycosphingolipid accumulation in type1 Gaucher’s disease ^20,21^, was approved for NPC treatment in Europe, Canada and Japan but denied FDA approval (although it is prescribed off label in the US). Miglustat may confer mild improvement in specific clinical symptoms but fails to prevent disease progression ^22,23^. 2-Hydroxy propyl beta cyclodextrin (HPBCD) is being investigated as an emerging therapy ^24,25^. It chelates cholesterol but does not cross the blood brain barrier^26^. Therefore, to treat neurological disease HPBCD must be directly delivered to the central nervous system (CNS; ^27,28^) which carries procedural risk of life-long therapy. Systemic delivery is needed to improve liver and other visceral organs but inexplicably, HPBCD is excluded from lung ^29,30^ and therefore of little benefit to end-stage advanced and frequently fatal bronchial disease. Arimoclomol is another emergent therapy for NPC ^31^, but its benefit for systemic disease especially in treatment of lung inflammatory disease remains unknown.

The TCF combines HPBCD, PEG and Vo in a defined formulation ^10^. Upon systemic injection, it increases the plasma exposure of Vo and boosts its delivery across the BBB to stimulate histone acetylation there. Although mice chronically treated for close to a year showed no metabolic toxicity ^10^, the effect of long term TCF exposure on key neurons, brain areas and overall progression of symptoms of neurodegeneration that mimic human disease, have not been assessed. Further, while HPBCD reduces systemic inflammation ^10,24,29^, it is excluded from lungs ^29,30^ and therefore whether the TCF promotes Vo delivery and therapeutic action in lungs remains unknown. Our findings on these points advance development of a new HDACi therapeutic strategy to treat NPC and other difficult-to-treat disorders that may benefit from epigenetic therapy

## Methods

### Materials

All fine chemicals including 2-hydroxypropyl-*β*-cyclodextrin (HPBCD) and polyethylene glycol 400 (PEG) were procured from Sigma (St Louis, MO, USA) unless otherwise indicated. Vorinostat was from Selleck Chemicals (Houston, TX, USA).

### Animals

*Npc1*^*nmf164*^ is a BALB/c strain carry a D1005G (A to G at cDNA bp 3163) mutation in the *Npc1* gene ^32^. A breeding pair of mutant mice were obtained from Robert P. Erickson, University of Arizona Health Sciences Center, Tucson, AZ, USA and is available at ‘The Jackson Laboratories.’ Homozygous mutants *(Npc1*^*nmf164*^) along with wild-type littermates *(Npc1*^*+/+*^), were generated in house by crossing heterozygous mutant *(Npc1*^*+/nmf164*^) males and females and genotyped as previously described ^10^. Wild type Balb/c mice were procured from Envigo (Indianapolis, IN, USA)

### Drug injection and organ harvest

The Triple combination formulation (TCF) is a mixture of vorinostat (50mg/kg), HPBCD (2000mg/Kg)), PEG 400 (45%) and DMSO (5%). Vorinostat (50mg/Kg) was made in 5% DMSO and 45% PEG. HPBCD was a 20% (w/v) solution and given dose of 2000mg/Kg. Detailed methodology on preparing drug solutions have been descried earlier ^10^. To enable comparative studies with prior regiments, all mice were given two doses of HPBCD at P7 and P15. From P21 onwards, mice received either HPBCD alone or TCF, as indicated. Vo was also initiated at P21. Injections were administered weekly through the intraperitoneal (i.p) route (and the injection volume used was 10ml/Kg body weight across all treatment groups). For lung histopathology, *Npc1*^*nmf164*^ mice were analyzed at 100-109 days of age. Long-term safety was assessed for 8-10 months either in *Npc1*^*+/nmf164*^ or commercially purchased wild type Balb/c mice. The animals were sacrificed by asphyxiation using CO2 and harvested organs were immersed fixed in 10% neutral buffered formalin (∼4% formaldehyde) for 24 hours at RT and subsequently stored in 70% alcohol until transfer to paraffin.

### Nissl and H&E staining

Paraffin-embedded sections (4–5 μm) were dewaxed in xylene and alcohol. For Nissl, brain sections were stained with acidified 0.1% cresyl violet for 7 min followed by two incubations in 95% ethanol of 5 min each. The sections were cleared in xylene and mounted in cytoseal XYL (Thermo Scientific, Kalamazoo, USA). H&E staining of brain and lung tissues was carried out by AML laboratories according to standard methods ^33^. Images were visualized with DPIan Apo 40x/1.00 oil immersion objective lens (Nikon) and captured on a Nikon Olympus microscope, using a Nikon digital DS-Fi1-U2 camera controlled by NIS-Elements F3.0 Nikon software (all from Nikon Instruments INC, Tokyo, Japan).

### Iba1 immunostaining of brain sections

Paraffin-embedded brain sections (4–5 μm) were dewaxed in xylene and alcohol. Iba1 antigen was retrieved by boiling the sections in acidic condition for 30 min. Blocking was done with 2% goat serum for 30 min at RT. Sections were incubated with anti-Iba1 (1:500, 019-19741, Wako Chemicals) overnight at 4 °C. FITC-conjugated secondary IgG antibodies (MP Biomedicals, Solon, OH, USA) were used at a dilution of 1:200. Nuclei were stained with DAPI (0.5μg/ml) and mounting was done using Vectashield (Vector laboratories). Sections were visualized with 40× oil-immersion objective lens (NA 1.35) and image collection was performed using an Olympus IX inverted fluorescence microscope and a Photometrix cooled CCD camera (CH350/LCCD) driven by DeltaVision software from Applied Precision (Seattle, WA, USA). DeltaVision software (softWoRx) was used to deconvolve these images. Images are single optical sections. Images were analyzed using ‘softWoRx’ or ‘ImageJ’ software (NIH, MD, USA).

### Quantification of Vo in lungs

*Npc1*^*+/nmf164*^ mice (age 6-7 weeks) were given intraperitoneal injections of either Vo (50mg/Kg in 45% PEG and 5% DMSO) or TCF (Vo 50mg/Kg + HPBCD, 2000mg/Kg + PEG, 45% + DMSO, 5%). At 30 min and 1 h post injection, mice were asphyxiated with CO2, blood was drawn by cardiac puncture and organs were perfused with 20 ml ice-cold PBS through the ventricle. Harvested lungs were cut into small pieces (4-6 mm^2^) and flash frozen in liquid nitrogen. The quantification of vorinostat was done by Metabolite Profiling Facility, Bindley Bioscience Center, Purdue University, IN, USA. The detailed methods are as described earlier ^10^. Briefly, the tissue was homogenized using a Precelly bead homogenizer system utilizing ceramic CK 14 beads. 2 ng of deuterated internal standard (d_5_-Vorinostat, Toronto Research Chemicals, Ontario, Canada) was added to lung homogenate prior to liquid extraction with acetonitrile. Prior to analysis, samples were reconstituted in 100 μL of 50% water/50% acetonitrile. An Agilent 1200 Rapid Resolution liquid chromatography (HPLC) system coupled to an Agilent 6460 series triple quadrupole mass spectrometer (MS/MS) was used to analyze vorinostat. The data were obtained in positive electrospray ionization (ESI) mode and quantitated by monitoring the following transitions: for Vorinostat, 265→232 with a collision energy of 5 V and for d5- Vorinostat, 270→237 with collision energy of 5 V.

## Results

### Assessment of chronic TCF-treatment in cerebellar and hippocampal regions as well as a neurobehavioral/cognitive disease score in mice

HDACs are important enzymes and their functions are required in brain development ^34-36^. In particular, HDAC3 knockdown blocked development of Purkinje neurons ^37^. It has therefore been hypothesized that long-term HDACi treatment may adversely affect the brain. However, we have reported that weekly administration of the TCF in *Npc1*^*nmf164*^ mice prevented loss of Purkinje cell neurons ^10^. Since Vorinostat from the TCF peaks at 30 minutes and is rapidly cleared from the brain and plasma ^10^, our findings suggested that epigenetic modulation associated with transient block of HDAC3 (as well as other HDACs) may be tolerated and benefit NPC-diseased animals.

But the effects of TCF on neurons in normal animal brain remain unaddressed. Since Purkinje are major neurons requiring HDAC function, we used them as a sentinel neuron for effects of extended TCF-treatment in healthy animals. We administered weekly TCF to heterozygous, healthy ‘control’ animals for 2-3 fold longer (240-300 days) than the 100 day-efficacy period in *Npci*^*nmf164*^ mice. As shown in Fig. 1a, H&E staining failed to show any change in histological features of Purkinje cells in the cerebellum. Nissl staining suggested they were intact neurons (Fig. 1b). Quantitative analyses of both H&E - and Nissl –staining confirmed that the TCF even on extended treatment did not lead death and loss of Purkinje cells (Fig. 1c). These data indicate that recurrent, short-lived exposure of low concentration of vorinostat does not cause neuronal loss, even as it is sufficient to trigger sustained epigenetic effects.

**Figure 1.**
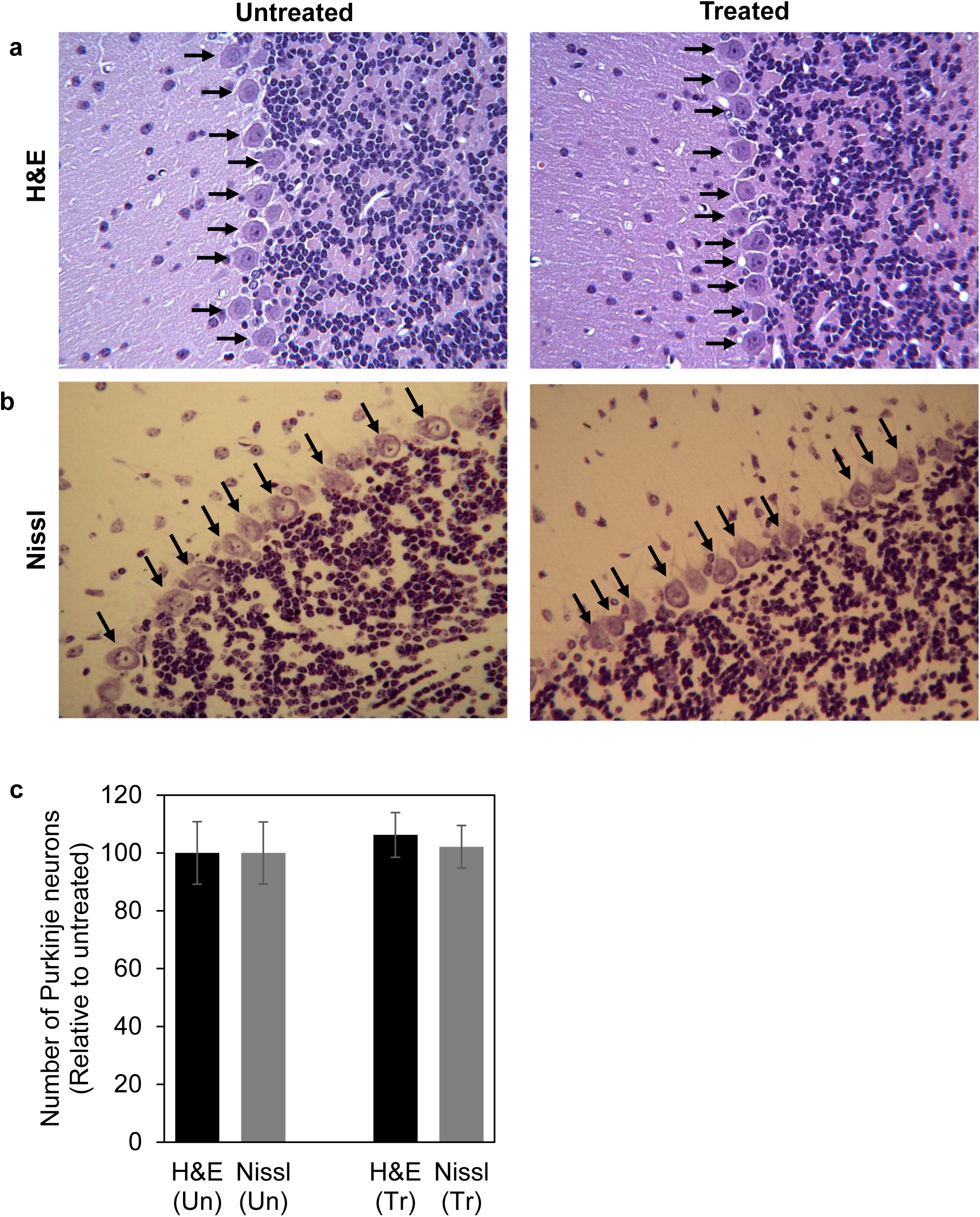
Analysis of Purkinje neurons in long-term TCF-treated mice by **(a)** H&E and **(b)** Nissl staining. Representative micrographs of cerebellum from untreated (108 days) and TCF-treated healthy *(Npc1*^*+/nmfI64*^, 225-265 days) mouse are shown. Purkinje neurons are shown by black arrows. The images shown are representative from four mice in each group. (c) Bar diagram is quantification of Purkinje neurons from same number of mice. Two sections per mouse were analyzed for counting. Numbers (mean±SD) are relative to untreated healthy mice. Un, untreated; Tr, treated.

Activation of microglial cells marks neuroinflammation, an early sign of neuronal dysfunction ^38,39^. We have previously shown that microglia stained with antibodies to the Iba inflammatory marker accumulate in the hippocampus of *Npc1*^*nmf164*^ mice ^10^. Further at ∼ 100 days after weekly TCF treatment, Iba staining is reduced suggesting that TCF can target inflammatory dysfunction in the hippocampus. Comparable H&E staining is seen in representative hippocampal regions from mice exposed to weekly TCF treatment for 225-265 days compared to untreated animals at 100-110 days (Fig. 2a) with each showing only a few resident microglial cells (Fig. 2b). Quantitative analysis showed no significant difference in the number of microglial cells untreated and chronically TCF-treated mice (Fig. 2c), suggesting that despite the fact that Vo is predicted to transcriptionally activate numerous target genes, the TCF does not induce an inflammatory response broadly damaging to neurons.

**Figure 2.**
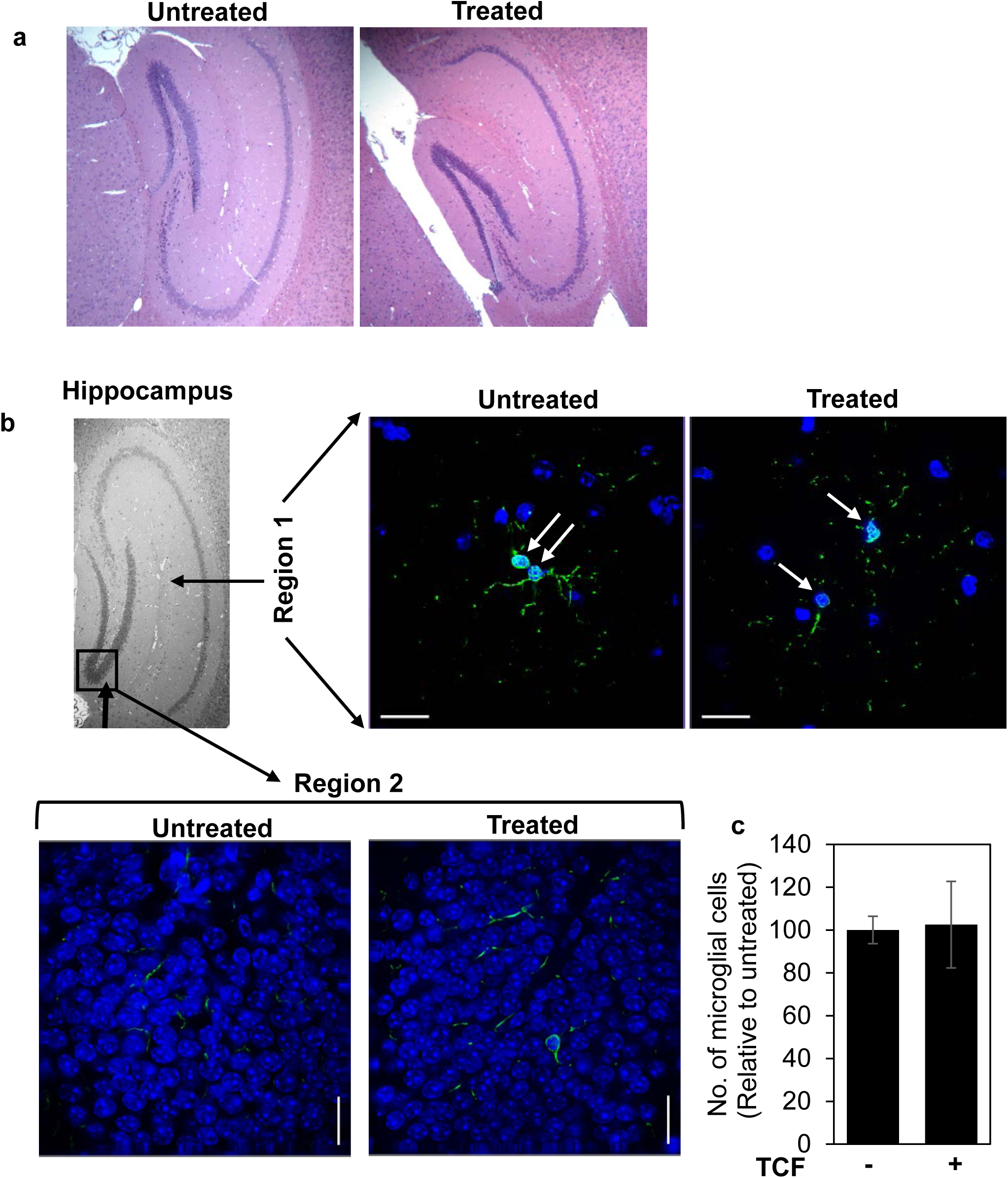
Safety assessment in the hippocampus of mice after chronic treatment with TCF for long-term. **(a)** Histological analysis using H&E staining **(b)** Analysis of neuroinflammation in long-term TCF-treated mice. Fluorescence microscopy detection of activated microglial cells (green, white arrows) with anti Iba-1 antibodies in the hippocampus (from the regions indicated) from untreated and TCF-treated healthy *(Npc1*^*+/nmf164*^). The images shown are representative from two untreated (age 108 and 109 days) and four TCF-treated (age 225-265 days) healthy mice, **(c)** Bar diagram is quantification of microglial cells from same number of mice. Two sections per mouse were studied. Scale bar, 25μm

Clinically NPC disease is defined by major and minor symptomatic domains, whose severity has been quantified to monitor the natural history of the disease ^40^. While awaiting plasma biomarkers ^41-44^ symptom scoring continues as an important index of progressive disease. We previously created a disease severity scale for murine NPC that captures major patient disease domain scored in a defined indicated range and whose sum provides the cumulate disease score, (with a maximal score of 13; ^10^). Because older healthy animals, particularly males, often displayed poor grooming and slight impairment in limb tone onwards of 100 days, a cumulative score of 3 or higher reliably flags onset of symptomatic disease. Untreated *Npc1*^*nmf164*^ mice progress to a score of 10-13 by 100 days, but TCF affords significant reduction to 4-5 when administered over the same period ^10^ to render functional benefit to major symptomatic domains of neurological disease that include ambulation, cognition, motor control and dysphagia. In contrast, the cumulative score remained below baseline in healthy wild type mice receiving chronic weekly administration of TCF (Fig. 3a, Supplementary Table S1; with indicated scores of 1-2 that also appear in untreated animals, as expected due to their poor grooming). There was also no change in animal weight (Fig. 3b) showing TCF-treatment did not impair overall nutrient consumption and utilization (which marks mid- and end-stage neurological disease ^10^).

**Figure 3.**
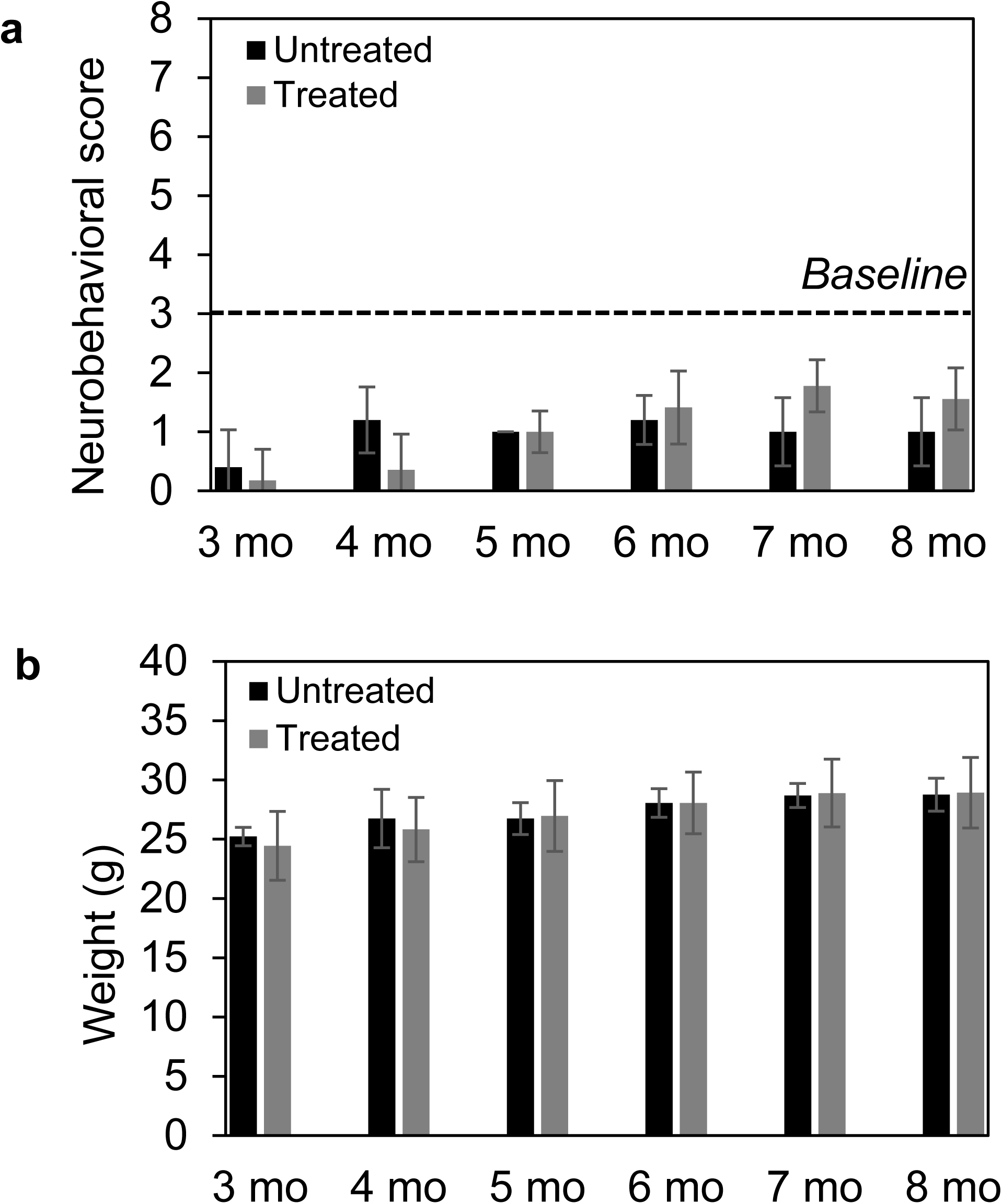
Assessment of chronic TCF treatment on neurobehavioral/cognitive disease score and body weight in wild type Balb/c mice. **(a)** Major neurobehavioral symptoms and (corresponding human disease domain) as follows: tremor (motor); gait (ambulation); grooming (cognition); body position (cognition and motor), limb tone (motor) and weight loss (dysphagia) were assessed on a scale of 0-2 except weight loss assessed as 0-3. The cumulative score is shown at indicated time points. Score of 3 or below is baseline. For 3-6 months (mo) untreated, n=15 (all males) and treated, n=17 (7 males and 10 females). For 7 and 8 months untreated, n=7 (all males) and treated, n=9 (4 males and 5 females), **(b)** Weight of mice in A. Data are mean ± SD. See also Supplementary Table S1.

### TCF increases Vo levels in lung

As previously reported the TCF is a triple combination formulation containing Vo, HPBCD and PEG ^10^. In prior work we found that 1 h after injection of TCF, Vo concentrations in mouse plasma were 3 fold higher compared to the levels observed when Vo was administered in PEG alone ^10^. Vo levels in the brain were also significantly boosted in TCF-injected mice ^10^. These data suggested that the HPBCD was a major contributor to the pharmacokinetic (PK) effect in plasma and brain. Further examination of brain, liver and spleen suggested the TCF could treat both neurological as well as systemic NPC disease in mice. However, since HPBCD is known to be excluded from lungs ^29,30^, it remained unclear whether the TCF increased exposure of Vo and/or benefit lung disease.

As shown in Fig. 4, animals injected with Vo in PEG alone showed a mean concentration of 3.2 ng/mg Vo in lungs at 30 minutes, which decreased to 1 ng/mg by 60 min. After TCF injection, Vo concentration reached 7.9 ng/mg at 30 min and then declined to 4.2 ng/mg at 60 min. These data suggested that the TCF boosted Vo entry into lungs, likely due to the (2.5-3 fold) plasma pharmacokinetic effect (previously reported in ^10^). Vo concentrations (of 4.2 ng/mg) detected at 60 min in TCF-treated animals were reduced by 45-50% reduction from levels seen at 30 min. Animals injected with Vo in PEG showed a 65-70% reduction over the same period, suggesting that in addition to boosting peak concentrations, the TCF may also slow down Vo clearance from lungs and both effects may increase levels and exposure of Vo in lungs.

**Figure 4.**
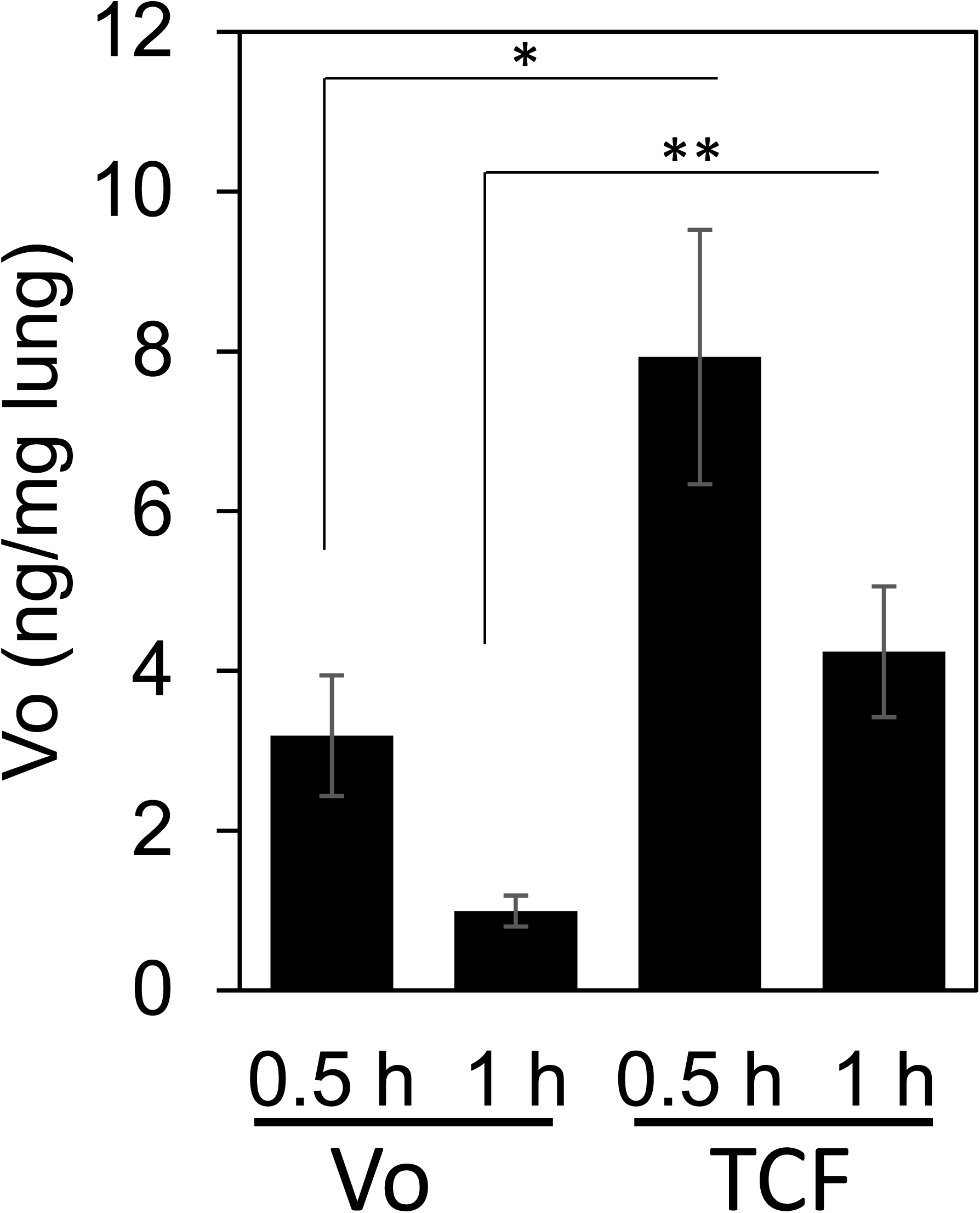
Increased lung concentration of Vo in TCF injected mice. *Npc1*^*+/nmf164*^ injected i.p. with Vo or TCF. At indicated times, animals sacrificed, perfused with PBS and Vo concentration in lungs were determined by mass spectrometry. n=5. h, hour. *p=0.02, TCF *vs* Vo 0.5 h, and * * p=0.014 TCF *vs* Vo, 1 h, two tailed Student’s *t* test.

### TCF reduces the accumulation of foamy macrophages in the lungs of *Npc1*^*nmf164*^ mice

Previous studies ^24,29,45^ have shown that the systemic delivery of HPBCD in NPC mice fails to alleviate lung disease (because HPBCD may not be able to reach the tissue). Since HPBCD is a major component of the TCF, this raised question on whether the formulation could alleviate lung disease even as it boosted Vo delivery to other organs. To test this, we undertook histochemical analysis of lungs from control and treated mice. Animals were examined at 100 days of age, since in prior work with the *Npc1*^*nmf164*^ mouse model; we have shown that this is a time of significant neurological disease, responsive to treatment by TCF. As shown in Fig. 5, H&E stained micrographs revealed accumulation of large number of foamy macrophages in the lungs of untreated *Npc1*^*nmf164*^ mice at 100 days. Semi-quantitative analysis showed TCF treatment significantly reduced the number of macrophages (Fig. 5). In contrast, administration of Vo alone or HPBCD continued to be associated with abundant macrophage accumulation (Fig. 5). Together these findings suggest that TCF-induced increase of Vo in lungs and reduced inflammation there.

**Figure 5.**
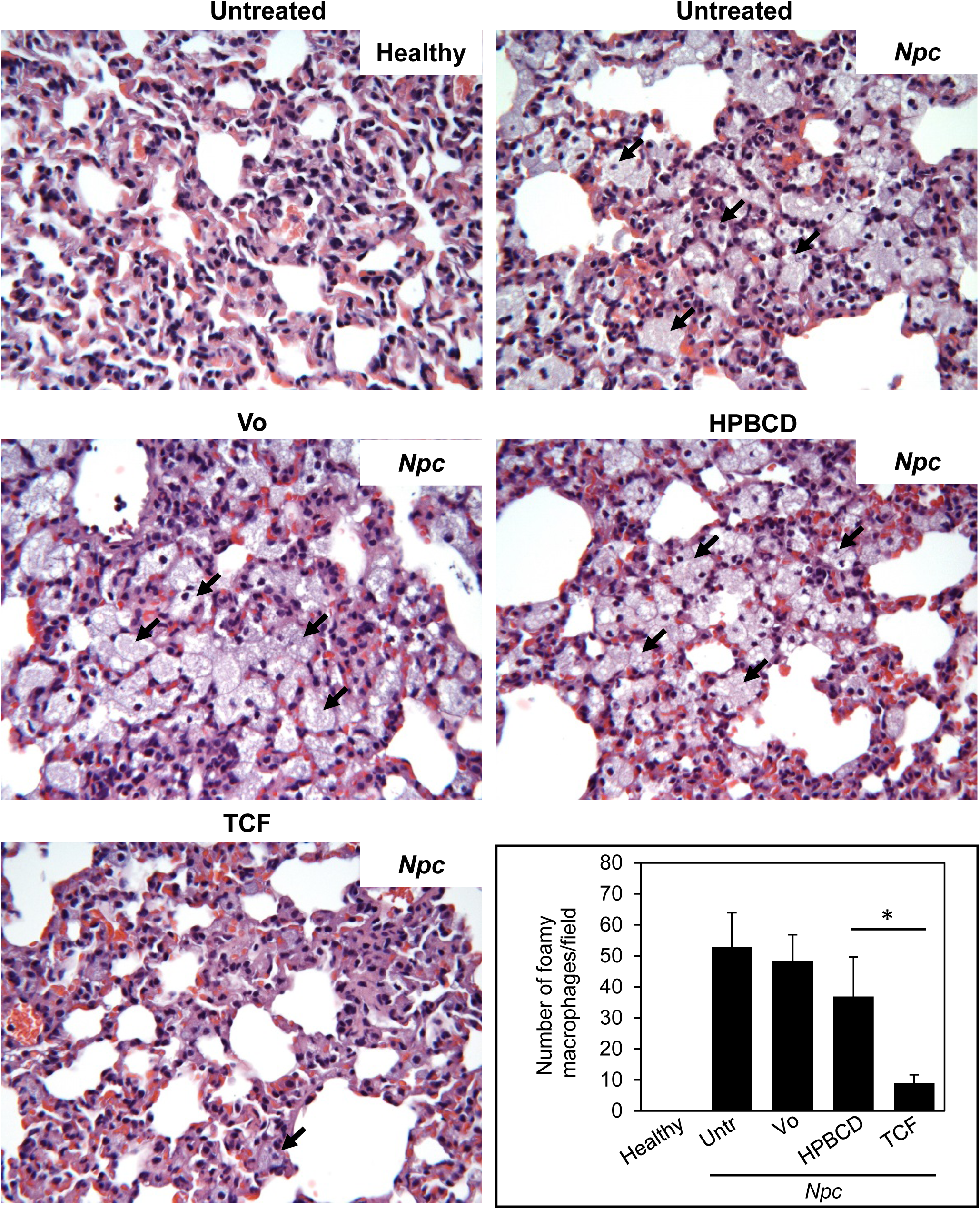
Efficacy of TCF in reducing the accumulation of foamy macrophages in the lungs of *Npc* mice. H&E stained micrographs showing foamy macrophages (indicated by black arrows) at 100 days of age in the lung of *Npc1*^*+/nmf164*^ (healthy control) and *Npc1*^*nmf164*^ *(Npc)* mutant mice treated as indicated. Foamy macrophages were abundant in untreated *Npc* mice. Treatment with Vo (vorinostat) or HPBCD had no effect whereas TCF treatment greatly reduced the accumulation of foamy macrophages. Images were taken with 40x objective lens and are representative of 4 mice in each group. Number of mice in each group=4. For quantitation, 10-15 random fields were analyzed per lung section. Untr, Untreated.* p=0.02, TCF vs HPBCD, twotailed Mann-Whitney test.

### Long-term chronic treatment with TCF shows no deleterious effect on lung histopathology in healthy control animals

Since our treatment analyses in *Npc1*^*nmf164*^ were undertaken at 100 days, the effects of extended TCF administration in control *Npc1*^*+/nmf164*^ type animals were assessed at 2-3 times longer periods (200-300 days). We previously reported that analysis of metabolic markers in the plasma failed to reveal toxicity in the liver and kidneys of mice treated once weekly with TCF after 200-300 days ^10^. Histological features of liver were also found to be normal ^10^. In Fig. 6, we show that lungs of *Npc1*^*+/nmf164*^ mice at 200-300 days show absence of tissue lesions, immune cell invasion or any abnormal pathology, as determined by H&E staining.

**Figure 6.**
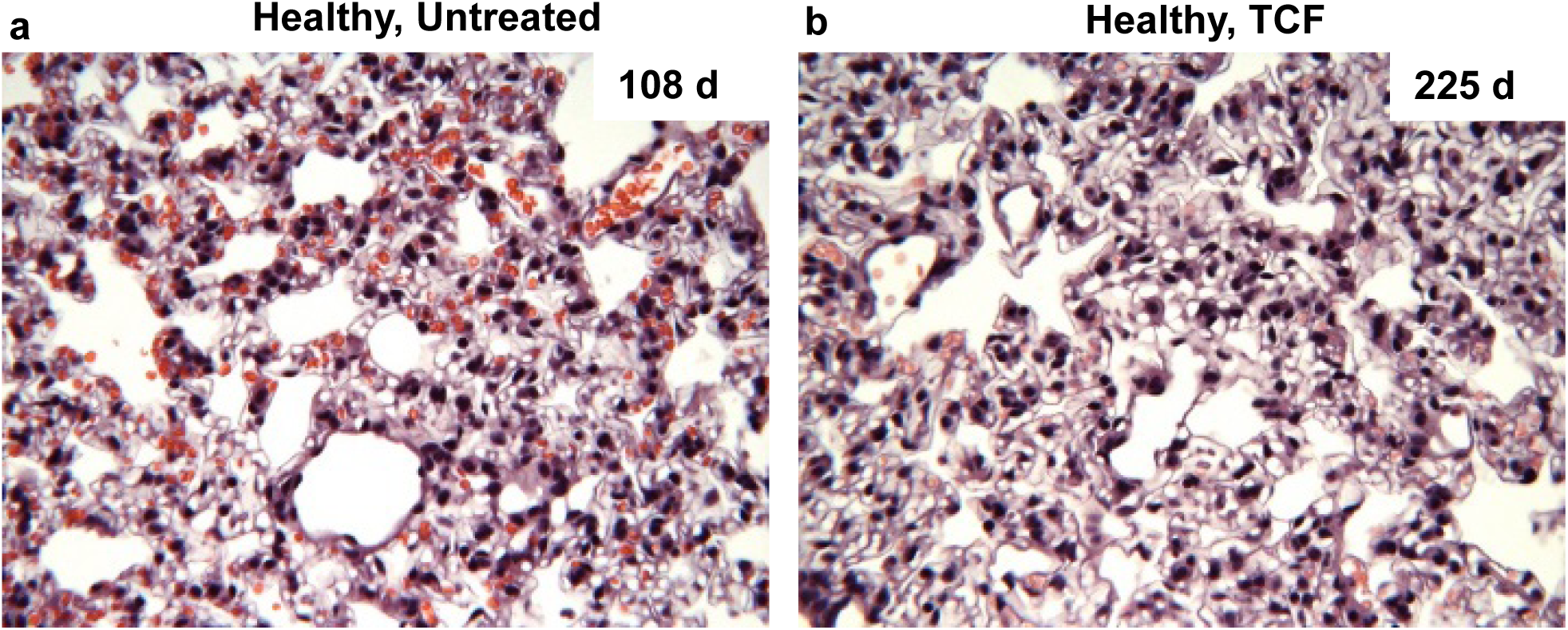
Histological analysis of lungs from long term TCF treated mice. Micrographs show **H&E** stained sections of lungs from (a) untreated and (b) TCF-treated healthy (*Npc1*^*+/nmf164*^) mouse at 108 and 225 days respectively. No signs of tissue lesions, immune cell invasion or abnormal pathology were seen in long term TCF-treated mice. The images shown are representative of four untreated (108-109 days) and four TCF-treated healthy mice (age 225-265 days). Images were taken with 40x objective lens

## Discussion

Concerns about intrinsic toxicity of HDACi are pertinent for both pan HDACi and inhibitors designed to target a given HDAC, since even a single HDAC can regulate hundreds of genes (and hence the value of synthesizing selective HDACi has been debated). Since neurological treatments may be long term, it is important to learn the effects of extended treatment periods well beyond when efficacy is detectable, especially in the brain. Our data in Figures 1-3 suggest that the TCF enables chronic administration of a therapeutically viable dose of broad spectrum HDACi with no detectable histological changes in key brain regions and neurocognitive/behavioral functions in mice. Purkinje neurons are major neurons that participate in motor control and learning. They can both emit and receive signals and function to regulate the entire cerebellum. Thus, maintenance of Purkinje cells provides a single read out for complex neuronal process in the cerebellum but also communication from the spinal cord and brain stem. Our data showing that the TCF helps preserve of Purkinje cells in the NPC disease model, suggests these cells are responsive to HDACi (likely due to elevation of NPC1 protein but possibly also by indirect mechanisms). Therefore, our finding that extended exposure to weekly TCF for 8 to 10 months had no effect on Purkinje neuron staining or count, suggests HDACi administration via the TCF is well tolerated in Purkinje-associated as well as overall cerebellar functions. Similarly, the hippocampus located in the cerebrum and a key region for learning and memory, shows no adverse structural and inflammatory effects despite extended TCF exposure. Although assessment of neurocognitive and behavioral scores do not yield quantitative tissue analyses, they indicate that TCF does not induce symptoms (and therefore processes) of neurodegeneration in wild type mice, even though it can delay appearance of these diseased processes in the NPC mouse model. Finally, findings that the TCF can boost delivery of vorinostat into lungs and reduce recruitment of macrophages into alveolar spaces, suggests that although HPBCD is excluded, vorinostat released from the TCF gains access lungs likely due the plasma exposure. Vo levels delivered to lungs are boosted to sufficiently reduce macrophage levels in *Npc1*^*nmf164*^ mice, which significantly expands the potential of the TCF in treating all organ systems expected to affect the progression of NPC. Extended TCF administration showed no ill-effects on lung pathology of normal mice

To conclude, extended TCF administration failed to induce adverse effects on metabolic parameters, brain and neurological functions as well as visceral organs including lung, although the TCF shows efficacy in all of these domains in the NPC mouse model. This may appear to be counterintuitive, since Vo is a broad acting HDACi at the transcriptional level with potential to target thousands of genes. However, proteomics studies suggest that changes may be limited to ∼200 targets in NPC diseased cells ^46^. Moreover, control healthy cells do not show major changes in proteome readouts in response to Vo ^46^. One explanation for this difference may be that mechanisms that restore normalcy in diseased cells are distinct from those that maintain homeostasis in normal cells.

Proteomic analyses of HDACi-induced changes in animal models of disease and health have yet to be undertaken. Moreover, in brain and lung, beneficial effects of Vo are detected at concentrations in the range of nanogram/mg which are substantially lower than those previously used to study Vo targets in cells and animals. It will be important to establish both transcriptional and proteomic targets of Vo in the 1-10 ng/mg range in both cells and animal organ systems. Finally, other than lung, HPBCD and PEG can access all organs in the body cavity and may act to modulate the effects of transcriptional/proteomic changes mediated by Vo primarily by lowering cholesterol and other lipid-related inflammatory pathways, to thereby further enhance long term tolerance of the TCF.

## Acknowledgements

We thank Brittany Coombs and Ashley VanAvermaete for technical assistance. The study was supported by the Parsons-Quinn Fund, University of Notre Dame.

### Author contributions

MSA: conceived and designed the study and its experiments, performed the experiments, analyzed data, wrote the paper. BC: conceived, designed and performed experimental studies on measurement of vorinostat in animal tissues. KH: conceived and designed the study and its experiments, analyzed the data and wrote the paper.

### Competing interests

KH holds equity interest in Ranedis Pharmaceuticals LLC. U.S. Provisional Application No. 62/011,553, entitled, “Formulation for the treatment of neurological diseases and cerebral injury” was filed on June 12, 2014”.

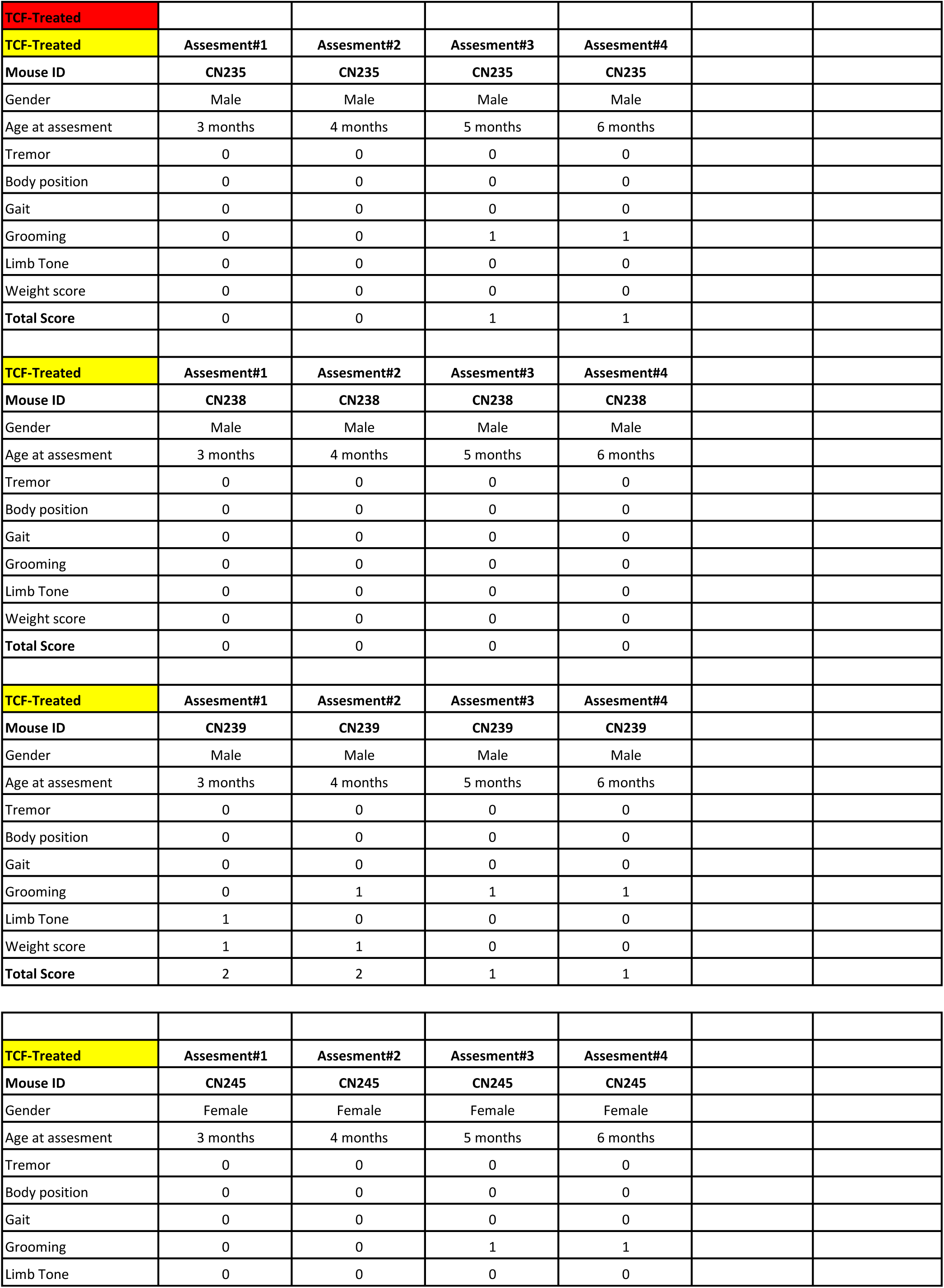
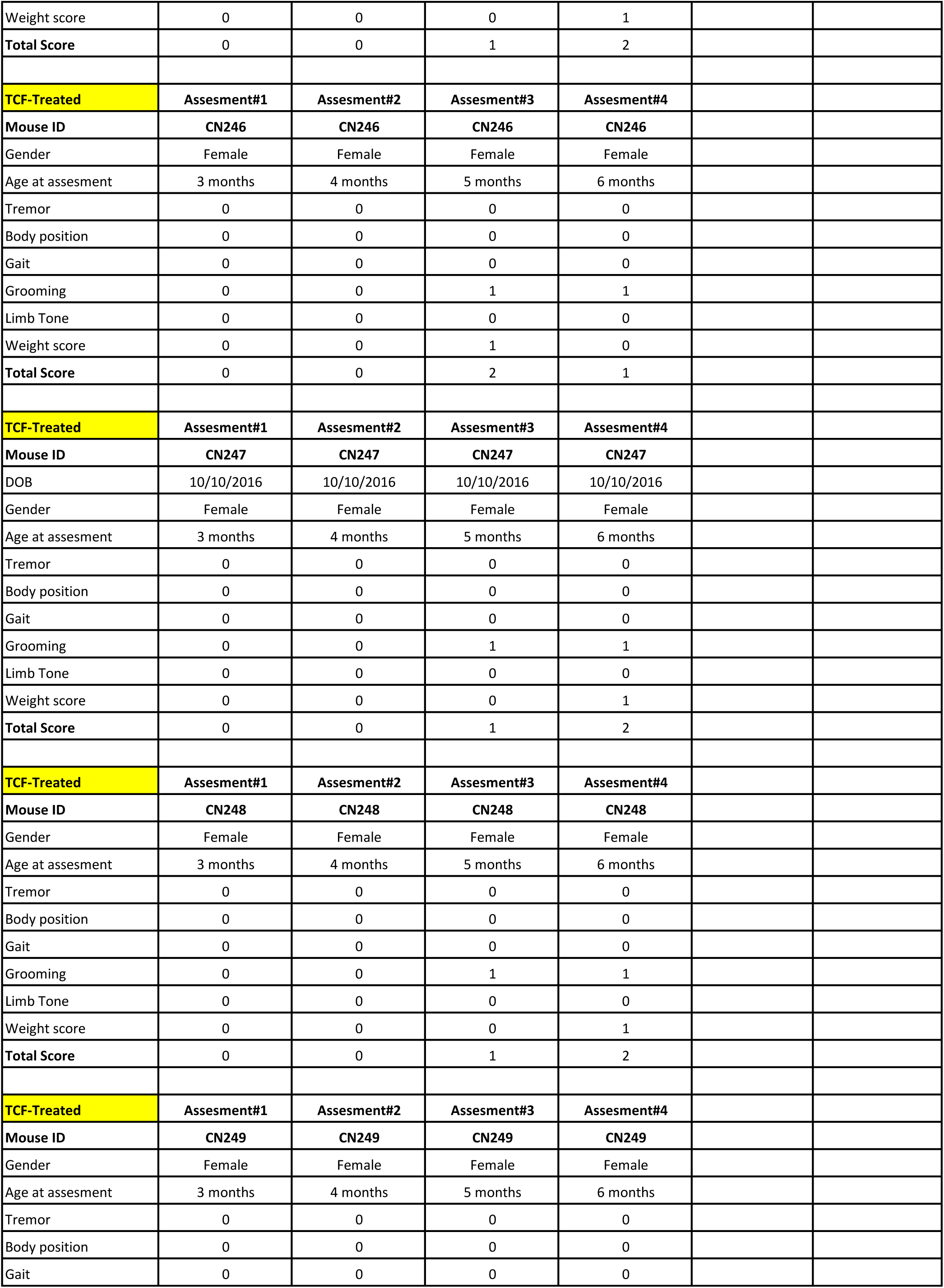
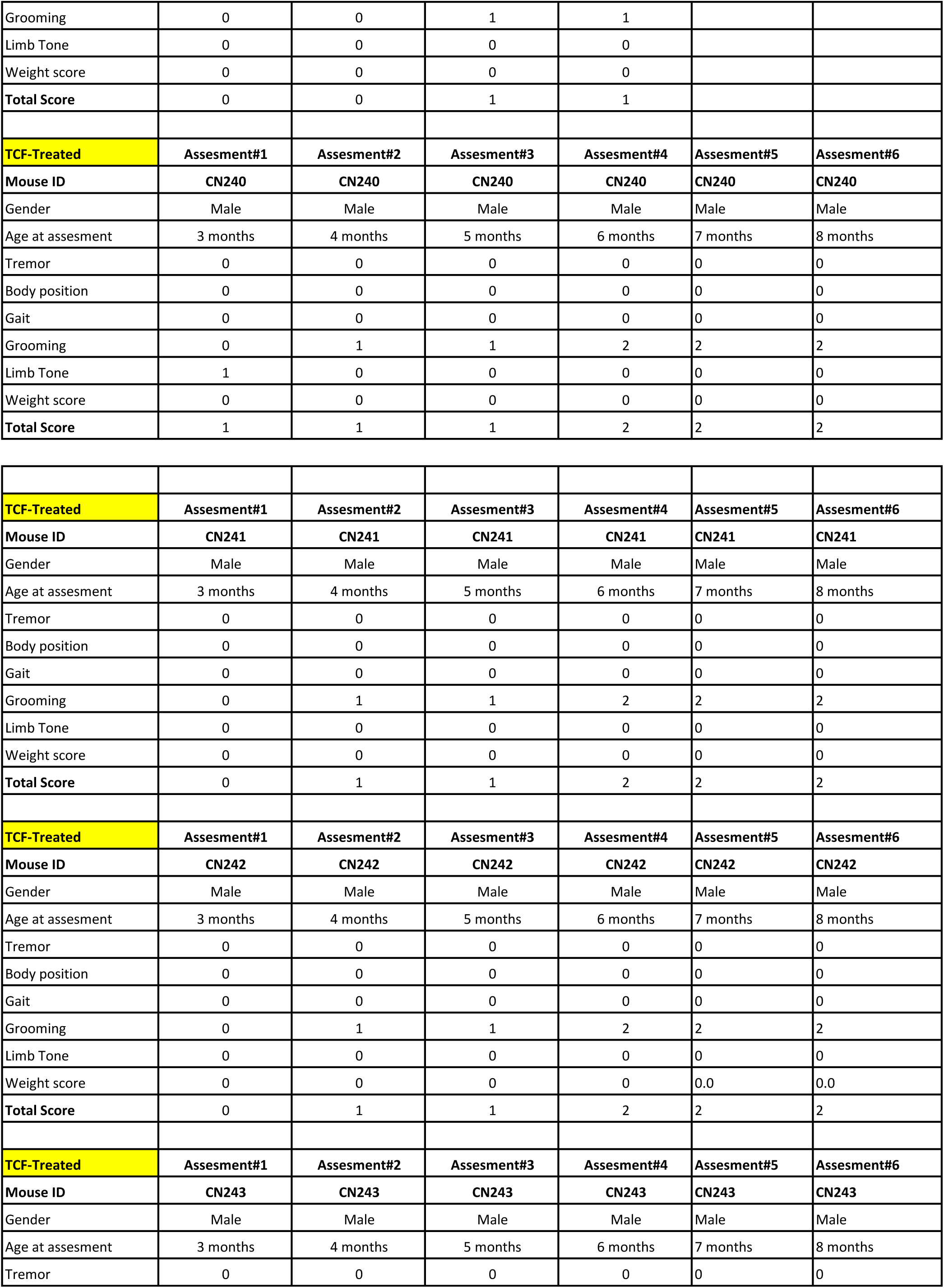
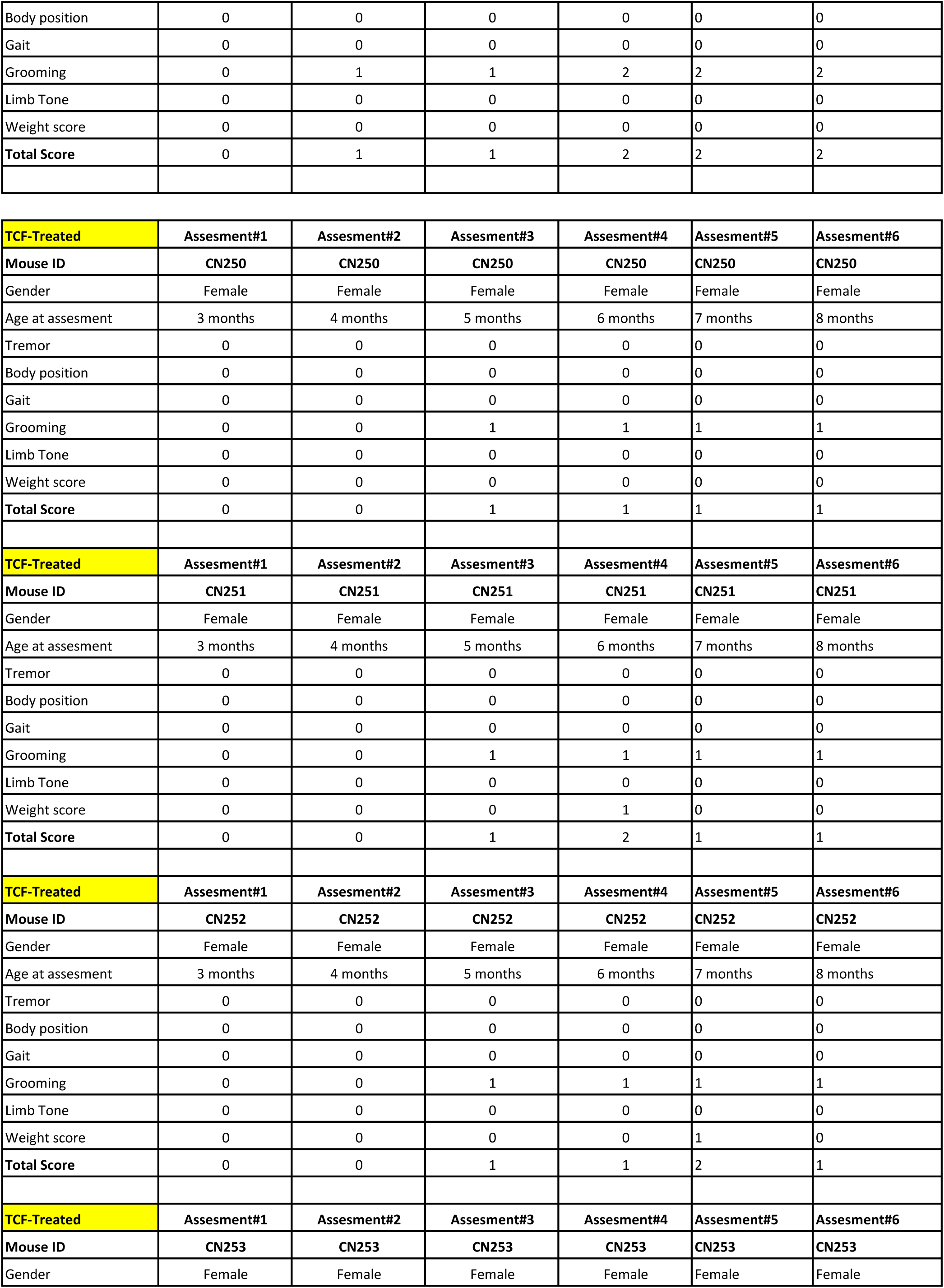
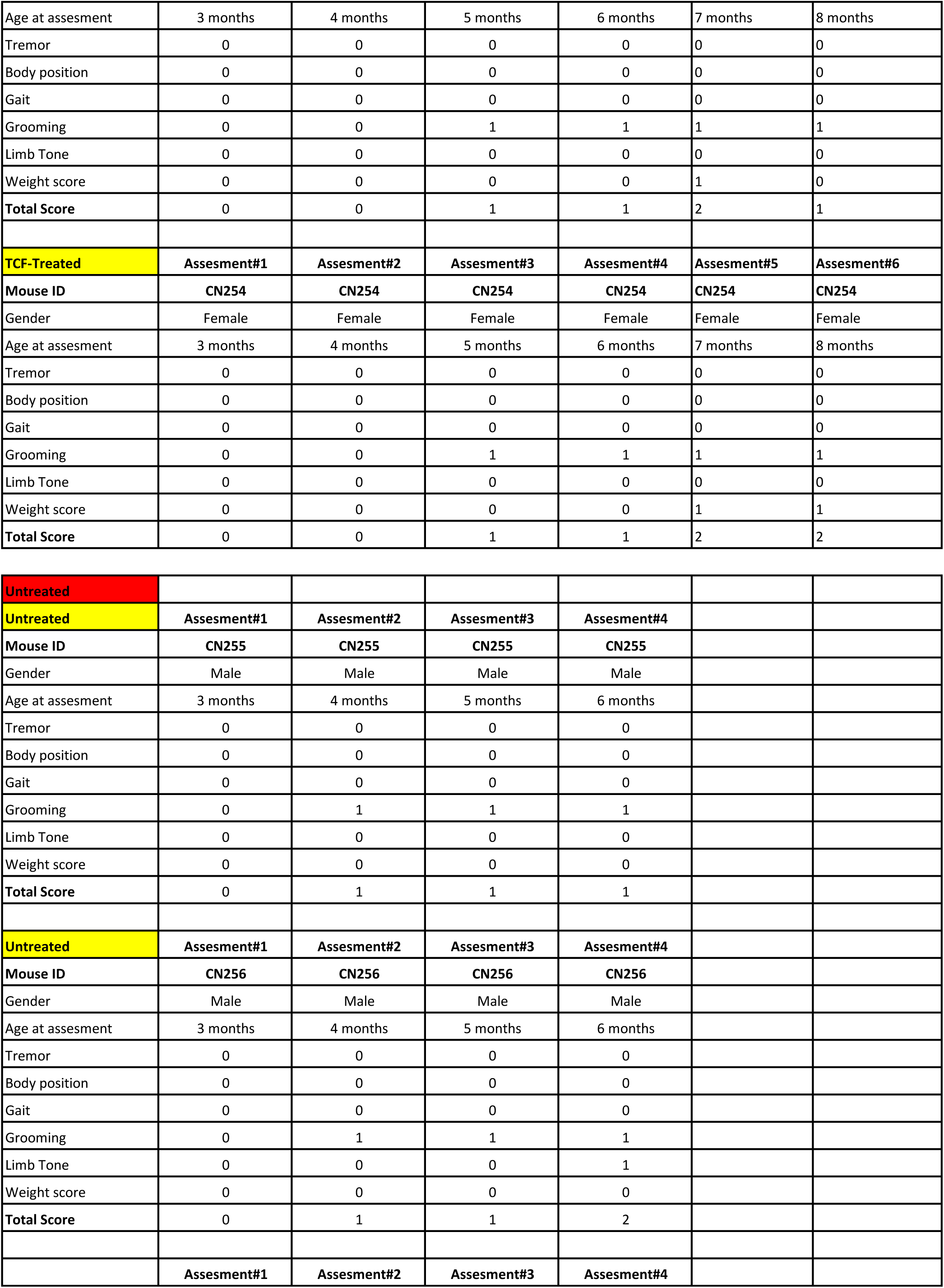
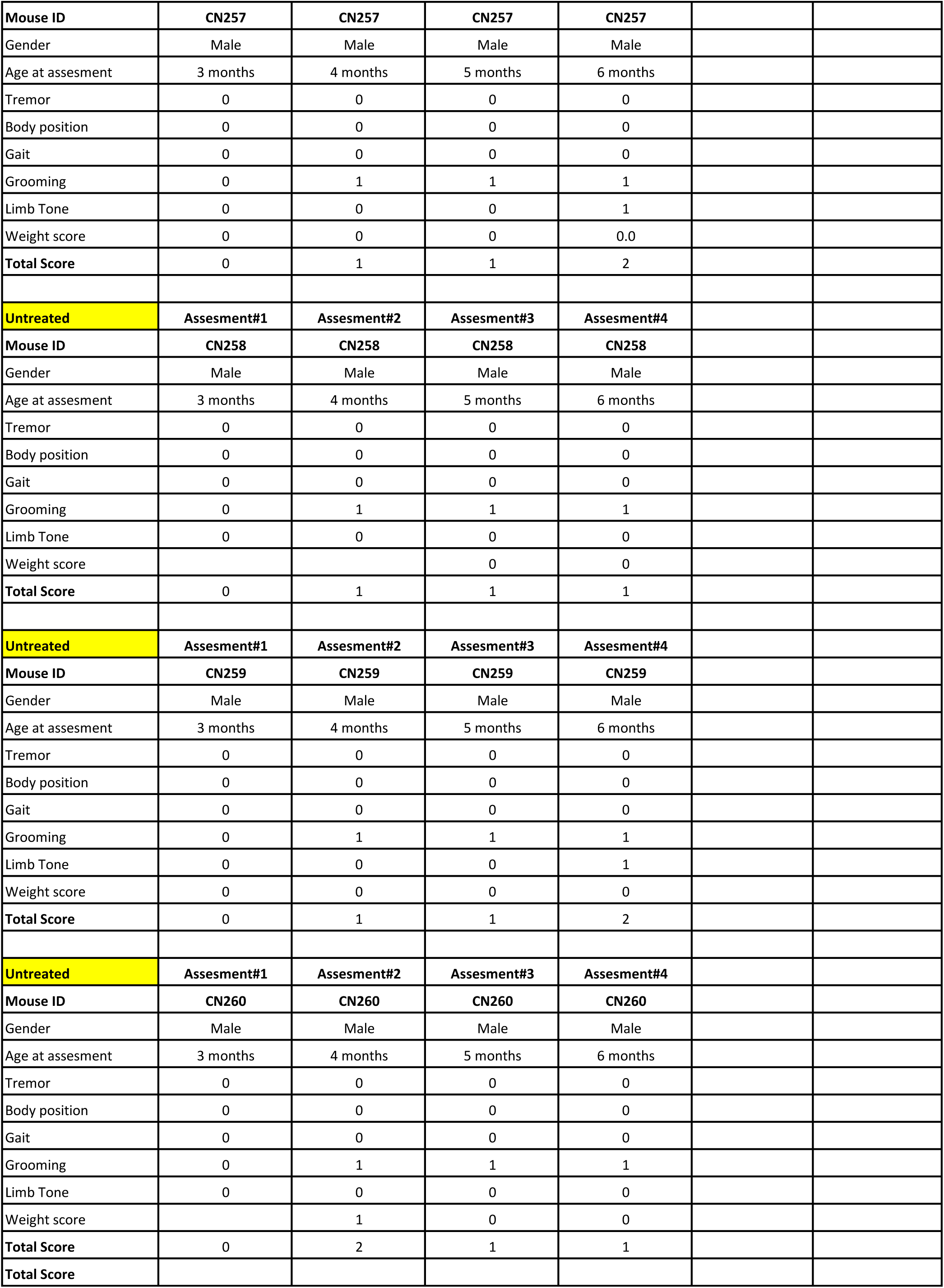
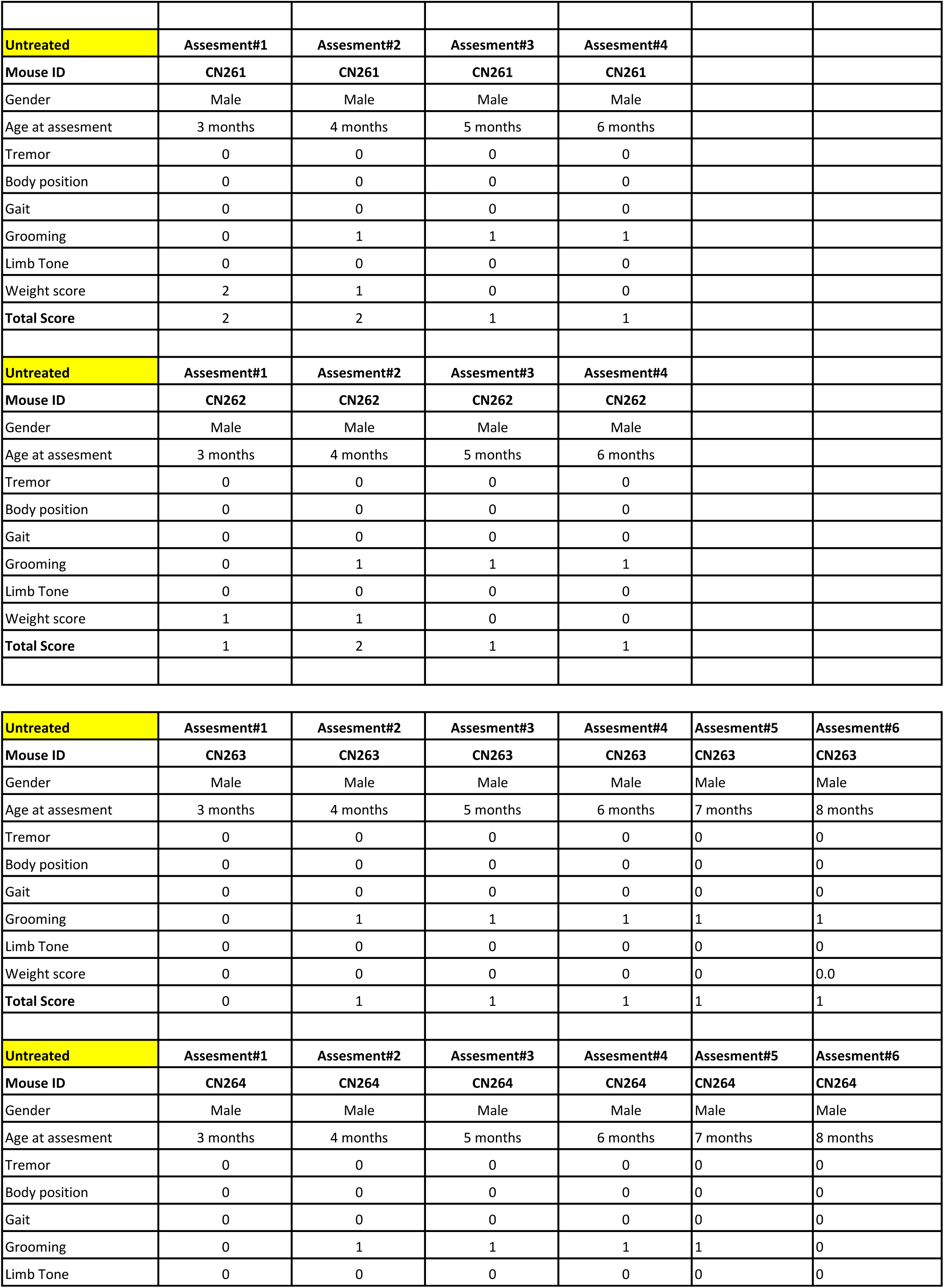
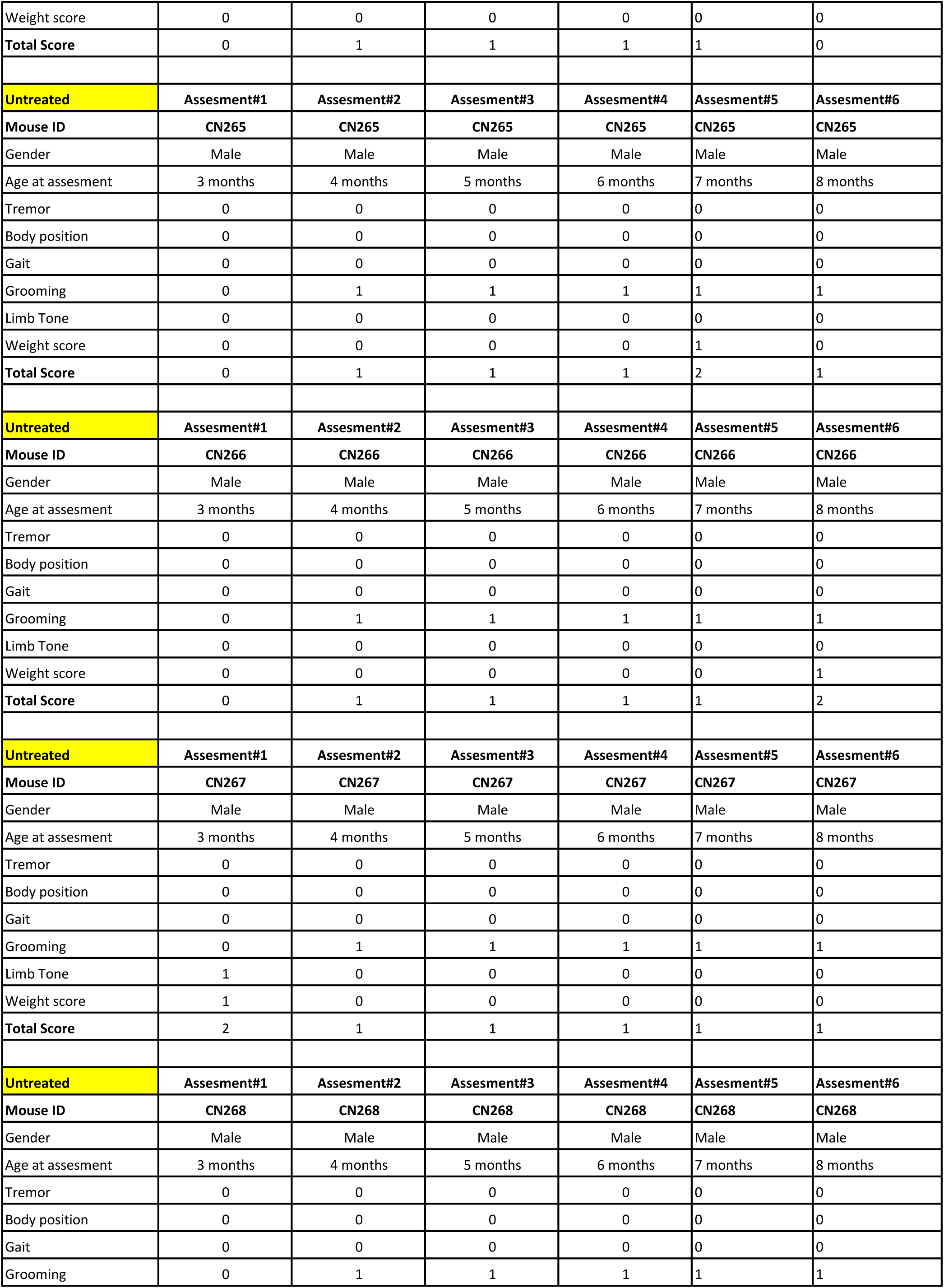
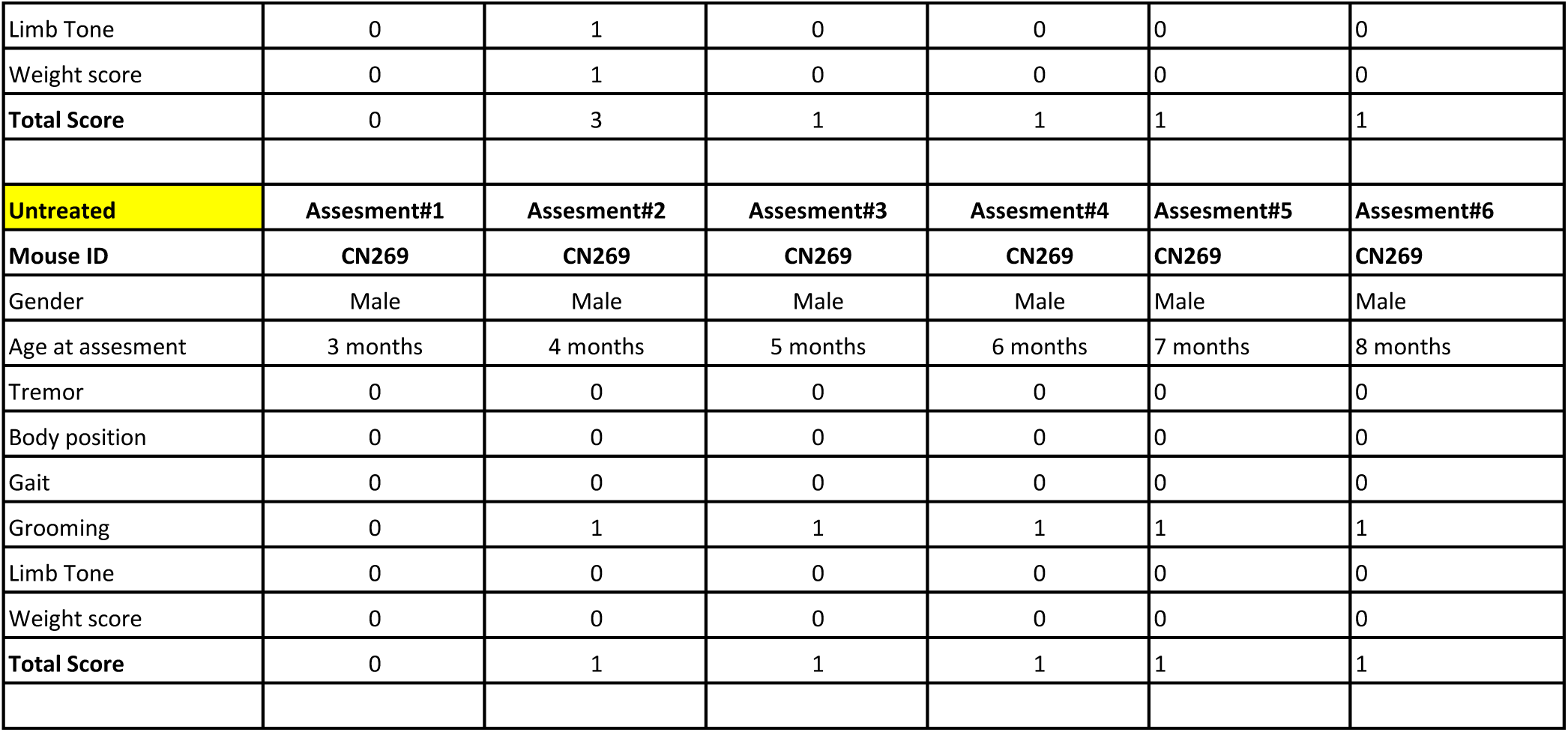

|  |  |  |  |  |
| --- | --- | --- | --- | --- |
| **TCF-Treated** |  |  |  |  |
| **TCF-Treated** | **Assesment#1** | **Assesment#2** | **Assesment#3** | **Assesment#4** |
| **Mouse ID** | **CN235** | **CN235** | **CN235** | **CN235** |
| Gender | Male | Male | Male | Male |
| Age at assesment | 3 months | 4 months | 5 months | 6 months |
| Tremor | 0 | 0 | 0 | 0 |
| Body position | 0 | 0 | 0 | 0 |
| Gait | 0 | 0 | 0 | 0 |
| Grooming | 0 | 0 | 1 | 1 |
| Limb Tone | 0 | 0 | 0 | 0 |
| Weight score | 0 | 0 | 0 | 0 |
| **Total Score** | **0** | **0** | **1** | **1** |
| **TCF-Treated** | **Assesment#1** | **Assesment#2** | **Assesment#3** | **Assesment#4** |
| **Mouse ID** | **CN238** | **CN238** | **CN238** | **CN238** |
| Gender | Male | Male | Male | Male |
| Age at assesment | 3 months | 4 months | 5 months | 6 months |
| Tremor | 0 | 0 | 0 | 0 |
| Body position | 0 | 0 | 0 | 0 |
| Gait | 0 | 0 | 0 | 0 |
| Grooming | 0 | 0 | 0 | 0 |
| Limb Tone | 0 | 0 | 0 | 0 |
| Weight score | 0 | 0 | 0 | 0 |
| **Total Score** | **0** | **0** | **0** | **0** |
| **TCF-Treated** | **Assesment#1** | **Assesment#2** | **Assesment#3** | **Assesment#4** |
| **Mouse ID** | **CN239** | **CN239** | **CN239** | **CN239** |
| Gender | Male | Male | Male | Male |
| Age at assesment | 3 months | 4 months | 5 months | 6 months |
| Tremor | 0 | 0 | 0 | 0 |
| Body position | 0 | 0 | 0 | 0 |
| Gait | 0 | 0 | 0 | 0 |
| Grooming | 0 | 1 | 1 | 1 |
| Limb Tone | 1 | 0 | 0 | 0 |
| Weight score | 1 | 1 | 0 | 0 |
| **Total Score** | **2** | **2** | **1** | **1** |
| **TCF-Treated** | **Assesment#1** | **Assesment#2** | **Assesment#3** | **Assesment#4** |
| **Mouse ID** | **CN245** | **CN245** | **CN245** | **CN245** |
| Gender | Female | Female | Female | Female |
| Age at assesment | 3 months | 4 months | 5 months | 6 months |
| Tremor | 0 | 0 | 0 | 0 |
| Body position | 0 | 0 | 0 | 0 |
| Gait | 0 | 0 | 0 | 0 |
| Grooming | 0 | 0 | 1 | 1 |
| Limb Tone | 0 | 0 | 0 | 0 |
| Weight score | 0 | 0 | 0 | 1 |
| **Total Score** | 0 | 0 | 1 | 2 |
| **TCF-Treated** | **Assesment#1** | **Assesment#2** | **Assesment#3** | **Assesment#4** |
| **Mouse ID** | **CN246** | **CN246** | **CN246** | **CN246** |
| Gender | Female | Female | Female | Female |
| Age at assesment | 3 months | 4 months | 5 months | 6 months |
| Tremor | 0 | 0 | 0 | 0 |
| Body position | 0 | 0 | 0 | 0 |
| Gait | 0 | 0 | 0 | 0 |
| Grooming | 0 | 0 | 1 | 1 |
| Limb Tone | 0 | 0 | 0 | 0 |
| Weight score | 0 | 0 | 1 | 0 |
| **Total Score** | 0 | 0 | 2 | 1 |
| **TCF-Treated** | **Assesment#1** | **Assesment#2** | **Assesment#3** | **Assesment#4** |
| **Mouse ID** | **CN247** | **CN247** | **CN247** | **CN247** |
| DOB | 10/10/2016 | 10/10/2016 | 10/10/2016 | 10/10/2016 |
| Gender | Female | Female | Female | Female |
| Age at assesment | 3 months | 4 months | 5 months | 6 months |
| Tremor | 0 | 0 | 0 | 0 |
| Body position | 0 | 0 | 0 | 0 |
| Gait | 0 | 0 | 0 | 0 |
| Grooming | 0 | 0 | 1 | 1 |
| Limb Tone | 0 | 0 | 0 | 0 |
| Weight score | 0 | 0 | 0 | 1 |
| **Total Score** | 0 | 0 | 1 | 2 |
| **TCF-Treated** | **Assesment#1** | **Assesment#2** | **Assesment#3** | **Assesment#4** |
| **Mouse ID** | **CN248** | **CN248** | **CN248** | **CN248** |
| Gender | Female | Female | Female | Female |
| Age at assesment | 3 months | 4 months | 5 months | 6 months |
| Tremor | 0 | 0 | 0 | 0 |
| Body position | 0 | 0 | 0 | 0 |
| Gait | 0 | 0 | 0 | 0 |
| Grooming | 0 | 0 | 1 | 1 |
| Limb Tone | 0 | 0 | 0 | 0 |
| Weight score | 0 | 0 | 0 | 1 |
| **Total Score** | 0 | 0 | 1 | 2 |
| **TCF-Treated** | **Assesment#1** | **Assesment#2** | **Assesment#3** | **Assesment#4** |
| **Mouse ID** | **CN249** | **CN249** | **CN249** | **CN249** |
| Gender | Female | Female | Female | Female |
| Age at assesment | 3 months | 4 months | 5 months | 6 months |
| Tremor | 0 | 0 | 0 | 0 |
| Body position | 0 | 0 | 0 | 0 |
| Gait | 0 | 0 | 0 | 0 |

|  |  |  |  |  |  |  |
| --- | --- | --- | --- | --- | --- | --- |
| Grooming | 0 | 0 | 1 | 1 |  |  |
| Limb Tone | 0 | 0 | 0 | 0 |  |  |
| Weight score | 0 | 0 | 0 | 0 |  |  |
| **Total Score** | 0 | 0 | 1 | 1 |  |  |
| **TCF-Treated** | **Assesment#1** | **Assesment#2** | **Assesment#3** | **Assesment#4** | **Assesment#5** | **Assesment#6** |
| **Mouse ID** | **CN240** | **CN240** | **CN240** | **CN240** | **CN240** | **CN240** |
| Gender | Male | Male | Male | Male | Male | Male |
| Age at assesment | 3 months | 4 months | 5 months | 6 months | 7 months | 8 months |
| Tremor | 0 | 0 | 0 | 0 | 0 | 0 |
| Body position | 0 | 0 | 0 | 0 | 0 | 0 |
| Gait | 0 | 0 | 0 | 0 | 0 | 0 |
| Grooming | 0 | 1 | 1 | 2 | 2 | 2 |
| Limb Tone | 1 | 0 | 0 | 0 | 0 | 0 |
| Weight score | 0 | 0 | 0 | 0 | 0 | 0 |
| **Total Score** | 1 | 1 | 1 | 2 | 2 | 2 |

|  |  |  |  |  |  |  |
| --- | --- | --- | --- | --- | --- | --- |
| **TCF-Treated** | **Assesment#1** | **Assesment#2** | **Assesment#3** | **Assesment#4** | **Assesment#5** | **Assesment#6** |
| **Mouse ID** | **CN241** | **CN241** | **CN241** | **CN241** | **CN241** | **CN241** |
| Gender | Male | Male | Male | Male | Male | Male |
| Age at assesment | 3 months | 4 months | 5 months | 6 months | 7 months | 8 months |
| Tremor | 0 | 0 | 0 | 0 | 0 | 0 |
| Body position | 0 | 0 | 0 | 0 | 0 | 0 |
| Gait | 0 | 0 | 0 | 0 | 0 | 0 |
| Grooming | 0 | 1 | 1 | 2 | 2 | 2 |
| Limb Tone | 0 | 0 | 0 | 0 | 0 | 0 |
| Weight score | 0 | 0 | 0 | 0 | 0 | 0 |
| **Total Score** | 0 | 1 | 1 | 2 | 2 | 2 |
| **TCF-Treated** | **Assesment#1** | **Assesment#2** | **Assesment#3** | **Assesment#4** | **Assesment#5** | **Assesment#6** |
| **Mouse ID** | **CN242** | **CN242** | **CN242** | **CN242** | **CN242** | **CN242** |
| Gender | Male | Male | Male | Male | Male | Male |
| Age at assesment | 3 months | 4 months | 5 months | 6 months | 7 months | 8 months |
| Tremor | 0 | 0 | 0 | 0 | 0 | 0 |
| Body position | 0 | 0 | 0 | 0 | 0 | 0 |
| Gait | 0 | 0 | 0 | 0 | 0 | 0 |
| Grooming | 0 | 1 | 1 | 2 | 2 | 2 |
| Limb Tone | 0 | 0 | 0 | 0 | 0 | 0 |
| Weight score | 0 | 0 | 0 | 0 | 0.0 | 0.0 |
| **Total Score** | 0 | 1 | 1 | 2 | 2 | 2 |
| **TCF-Treated** | **Assesment#1** | **Assesment#2** | **Assesment#3** | **Assesment#4** | **Assesment#5** | **Assesment#6** |
| **Mouse ID** | **CN243** | **CN243** | **CN243** | **CN243** | **CN243** | **CN243** |
| Gender | Male | Male | Male | Male | Male | Male |
| Age at assesment | 3 months | 4 months | 5 months | 6 months | 7 months | 8 months |
| Tremor | 0 | 0 | 0 | 0 | 0 | 0 |

|  |  |  |  |  |  |  |
| --- | --- | --- | --- | --- | --- | --- |
| Body position | 0 | 0 | 0 | 0 | 0 | 0 |
| Gait | 0 | 0 | 0 | 0 | 0 | 0 |
| Grooming | 0 | 1 | 1 | 2 | 2 | 2 |
| Limb Tone | 0 | 0 | 0 | 0 | 0 | 0 |
| Weight score | 0 | 0 | 0 | 0 | 0 | 0 |
| **Total Score** | 0 | 1 | 1 | 2 | 2 | 2 |

|  |  |  |  |  |  |  |
| --- | --- | --- | --- | --- | --- | --- |
| **TCF-Treated** | **Assesment#1** | **Assesment#2** | **Assesment#3** | **Assesment#4** | **Assesment#5** | **Assesment#6** |
| **Mouse ID** | **CN250** | **CN250** | **CN250** | **CN250** | **CN250** | **CN250** |
| Gender | Female | Female | Female | Female | Female | Female |
| Age at assesment | 3 months | 4 months | 5 months | 6 months | 7 months | 8 months |
| Tremor | 0 | 0 | 0 | 0 | 0 | 0 |
| Body position | 0 | 0 | 0 | 0 | 0 | 0 |
| Gait | 0 | 0 | 0 | 0 | 0 | 0 |
| Grooming | 0 | 0 | 1 | 1 | 1 | 1 |
| Limb Tone | 0 | 0 | 0 | 0 | 0 | 0 |
| Weight score | 0 | 0 | 0 | 0 | 0 | 0 |
| **Total Score** | 0 | 0 | 1 | 1 | 1 | 1 |

|  |  |  |  |  |  |  |
| --- | --- | --- | --- | --- | --- | --- |
| **TCF-Treated** | **Assesment#1** | **Assesment#2** | **Assesment#3** | **Assesment#4** | **Assesment#5** | **Assesment#6** |
| **Mouse ID** | **CN251** | **CN251** | **CN251** | **CN251** | **CN251** | **CN251** |
| Gender | Female | Female | Female | Female | Female | Female |
| Age at assesment | 3 months | 4 months | 5 months | 6 months | 7 months | 8 months |
| Tremor | 0 | 0 | 0 | 0 | 0 | 0 |
| Body position | 0 | 0 | 0 | 0 | 0 | 0 |
| Gait | 0 | 0 | 0 | 0 | 0 | 0 |
| Grooming | 0 | 0 | 1 | 1 | 1 | 1 |
| Limb Tone | 0 | 0 | 0 | 0 | 0 | 0 |
| Weight score | 0 | 0 | 0 | 1 | 0 | 0 |
| **Total Score** | 0 | 0 | 1 | 2 | 1 | 1 |

|  |  |  |  |  |  |  |
| --- | --- | --- | --- | --- | --- | --- |
| **TCF-Treated** | **Assesment#1** | **Assesment#2** | **Assesment#3** | **Assesment#4** | **Assesment#5** | **Assesment#6** |
| **Mouse ID** | **CN252** | **CN252** | **CN252** | **CN252** | **CN252** | **CN252** |
| Gender | Female | Female | Female | Female | Female | Female |
| Age at assesment | 3 months | 4 months | 5 months | 6 months | 7 months | 8 months |
| Tremor | 0 | 0 | 0 | 0 | 0 | 0 |
| Body position | 0 | 0 | 0 | 0 | 0 | 0 |
| Gait | 0 | 0 | 0 | 0 | 0 | 0 |
| Grooming | 0 | 0 | 1 | 1 | 1 | 1 |
| Limb Tone | 0 | 0 | 0 | 0 | 0 | 0 |
| Weight score | 0 | 0 | 0 | 0 | 1 | 0 |
| **Total Score** | 0 | 0 | 1 | 1 | 2 | 1 |

|  |  |  |  |  |  |  |
| --- | --- | --- | --- | --- | --- | --- |
| **TCF-Treated** | **Assesment#1** | **Assesment#2** | **Assesment#3** | **Assesment#4** | **Assesment#5** | **Assesment#6** |
| **Mouse ID** | **CN253** | **CN253** | **CN253** | **CN253** | **CN253** | **CN253** |
| Gender | Female | Female | Female | Female | Female | Female |

| Age at assesment | 3 months | 4 months | 5 months | 6 months | 7 months | 8 months |
| --- | --- | --- | --- | --- | --- | --- |
| Tremor | 0 | 0 | 0 | 0 | 0 | 0 |
| Body position | 0 | 0 | 0 | 0 | 0 | 0 |
| Gait | 0 | 0 | 0 | 0 | 0 | 0 |
| Grooming | 0 | 0 | 1 | 1 | 1 | 1 |
| Limb Tone | 0 | 0 | 0 | 0 | 0 | 0 |
| Weight score | 0 | 0 | 0 | 0 | 1 | 0 |
| **Total Score** | 0 | 0 | 1 | 1 | 2 | 1 |
| **TCF-Treated** | **Assesment#1** | **Assesment#2** | **Assesment#3** | **Assesment#4** | **Assesment#5** | **Assesment#6** |
| **Mouse ID** | **CN254** | **CN254** | **CN254** | **CN254** | **CN254** | **CN254** |
| Gender | Female | Female | Female | Female | Female | Female |
| Age at assesment | 3 months | 4 months | 5 months | 6 months | 7 months | 8 months |
| Tremor | 0 | 0 | 0 | 0 | 0 | 0 |
| Body position | 0 | 0 | 0 | 0 | 0 | 0 |
| Gait | 0 | 0 | 0 | 0 | 0 | 0 |
| Grooming | 0 | 0 | 1 | 1 | 1 | 1 |
| Limb Tone | 0 | 0 | 0 | 0 | 0 | 0 |
| Weight score | 0 | 0 | 0 | 0 | 1 | 1 |
| **Total Score** | 0 | 0 | 1 | 1 | 2 | 2 |
| **Untreated** |  |  |  |  |  |  |
| **Untreated** | **Assesment#1** | **Assesment#2** | **Assesment#3** | **Assesment#4** |  |  |
| **Mouse ID** | **CN255** | **CN255** | **CN255** | **CN255** |  |  |
| Gender | Male | Male | Male | Male |  |  |
| Age at assesment | 3 months | 4 months | 5 months | 6 months |  |  |
| Tremor | 0 | 0 | 0 | 0 |  |  |
| Body position | 0 | 0 | 0 | 0 |  |  |
| Gait | 0 | 0 | 0 | 0 |  |  |
| Grooming | 0 | 1 | 1 | 1 |  |  |
| Limb Tone | 0 | 0 | 0 | 0 |  |  |
| Weight score | 0 | 0 | 0 | 0 |  |  |
| **Total Score** | 0 | 1 | 1 | 1 |  |  |
| **Untreated** | **Assesment#1** | **Assesment#2** | **Assesment#3** | **Assesment#4** |  |  |
| **Mouse ID** | **CN256** | **CN256** | **CN256** | **CN256** |  |  |
| Gender | Male | Male | Male | Male |  |  |
| Age at assesment | 3 months | 4 months | 5 months | 6 months |  |  |
| Tremor | 0 | 0 | 0 | 0 |  |  |
| Body position | 0 | 0 | 0 | 0 |  |  |
| Gait | 0 | 0 | 0 | 0 |  |  |
| Grooming | 0 | 1 | 1 | 1 |  |  |
| Limb Tone | 0 | 0 | 0 | 1 |  |  |
| Weight score | 0 | 0 | 0 | 0 |  |  |
| **Total Score** | 0 | 1 | 1 | 2 |  |  |
|  | **Assesment#1** | **Assesment#2** | **Assesment#3** | **Assesment#4** |  |  |
| **Mouse ID** | **CN257** | **CN257** | **CN257** | **CN257** |  |  |
| Gender | Male | Male | Male | Male |  |  |
| Age at assesment | 3 months | 4 months | 5 months | 6 months |  |  |
| Tremor | 0 | 0 | 0 | 0 |  |  |
| Body position | 0 | 0 | 0 | 0 |  |  |
| Gait | 0 | 0 | 0 | 0 |  |  |
| Grooming | 0 | 1 | 1 | 1 |  |  |
| Limb Tone | 0 | 0 | 0 | 1 |  |  |
| Weight score | 0 | 0 | 0 | 0.0 |  |  |
| **Total Score** | 0 | 1 | 1 | 2 |  |  |
| **Untreated** | **Assesment#1** | **Assesment#2** | **Assesment#3** | **Assesment#4** |  |  |
| **Mouse ID** | **CN258** | **CN258** | **CN258** | **CN258** |  |  |
| Gender | Male | Male | Male | Male |  |  |
| Age at assesment | 3 months | 4 months | 5 months | 6 months |  |  |
| Tremor | 0 | 0 | 0 | 0 |  |  |
| Body position | 0 | 0 | 0 | 0 |  |  |
| Gait | 0 | 0 | 0 | 0 |  |  |
| Grooming | 0 | 1 | 1 | 1 |  |  |
| Limb Tone | 0 | 0 | 0 | 0 |  |  |
| Weight score |  |  | 0 | 0 |  |  |
| **Total Score** | 0 | 1 | 1 | 1 |  |  |
| **Untreated** | **Assesment#1** | **Assesment#2** | **Assesment#3** | **Assesment#4** |  |  |
| **Mouse ID** | **CN259** | **CN259** | **CN259** | **CN259** |  |  |
| Gender | Male | Male | Male | Male |  |  |
| Age at assesment | 3 months | 4 months | 5 months | 6 months |  |  |
| Tremor | 0 | 0 | 0 | 0 |  |  |
| Body position | 0 | 0 | 0 | 0 |  |  |
| Gait | 0 | 0 | 0 | 0 |  |  |
| Grooming | 0 | 1 | 1 | 1 |  |  |
| Limb Tone | 0 | 0 | 0 | 1 |  |  |
| Weight score | 0 | 0 | 0 | 0 |  |  |
| **Total Score** | 0 | 1 | 1 | 2 |  |  |
| **Untreated** | **Assesment#1** | **Assesment#2** | **Assesment#3** | **Assesment#4** |  |  |
| **Mouse ID** | **CN260** | **CN260** | **CN260** | **CN260** |  |  |
| Gender | Male | Male | Male | Male |  |  |
| Age at assesment | 3 months | 4 months | 5 months | 6 months |  |  |
| Tremor | 0 | 0 | 0 | 0 |  |  |
| Body position | 0 | 0 | 0 | 0 |  |  |
| Gait | 0 | 0 | 0 | 0 |  |  |
| Grooming | 0 | 1 | 1 | 1 |  |  |
| Limb Tone | 0 | 0 | 0 | 0 |  |  |
| Weight score |  | 1 | 0 | 0 |  |  |
| **Total Score** | 0 | 2 | 1 | 1 |  |  |
| **Total Score** |  |  |  |  |  |  |
| **Untreated** | **Assesment#1** | **Assesment#2** | **Assesment#3** | **Assesment#4** |  |  |
| **Mouse ID** | **CN261** | **CN261** | **CN261** | **CN261** |  |  |
| Gender | Male | Male | Male | Male |  |  |
| Age at assesment | 3 months | 4 months | 5 months | 6 months |  |  |
| Tremor | 0 | 0 | 0 | 0 |  |  |
| Body position | 0 | 0 | 0 | 0 |  |  |
| Gait | 0 | 0 | 0 | 0 |  |  |
| Grooming | 0 | 1 | 1 | 1 |  |  |
| Limb Tone | 0 | 0 | 0 | 0 |  |  |
| Weight score | 2 | 1 | 0 | 0 |  |  |
| **Total Score** | **2** | **2** | **1** | **1** |  |  |
| **Untreated** | **Assesment#1** | **Assesment#2** | **Assesment#3** | **Assesment#4** |  |  |
| **Mouse ID** | **CN262** | **CN262** | **CN262** | **CN262** |  |  |
| Gender | Male | Male | Male | Male |  |  |
| Age at assesment | 3 months | 4 months | 5 months | 6 months |  |  |
| Tremor | 0 | 0 | 0 | 0 |  |  |
| Body position | 0 | 0 | 0 | 0 |  |  |
| Gait | 0 | 0 | 0 | 0 |  |  |
| Grooming | 0 | 1 | 1 | 1 |  |  |
| Limb Tone | 0 | 0 | 0 | 0 |  |  |
| Weight score | 1 | 1 | 0 | 0 |  |  |
| **Total Score** | **1** | **2** | **1** | **1** |  |  |
| **Untreated** | **Assesment#1** | **Assesment#2** | **Assesment#3** | **Assesment#4** | **Assesment#5** | **Assesment#6** |
| **Mouse ID** | **CN263** | **CN263** | **CN263** | **CN263** | **CN263** | **CN263** |
| Gender | Male | Male | Male | Male | Male | Male |
| Age at assesment | 3 months | 4 months | 5 months | 6 months | 7 months | 8 months |
| Tremor | 0 | 0 | 0 | 0 | 0 | 0 |
| Body position | 0 | 0 | 0 | 0 | 0 | 0 |
| Gait | 0 | 0 | 0 | 0 | 0 | 0 |
| Grooming | 0 | 1 | 1 | 1 | 1 | 1 |
| Limb Tone | 0 | 0 | 0 | 0 | 0 | 0 |
| Weight score | 0 | 0 | 0 | 0 | 0 | 0.0 |
| **Total Score** | **0** | **1** | **1** | **1** | **1** | **1** |
| **Untreated** | **Assesment#1** | **Assesment#2** | **Assesment#3** | **Assesment#4** | **Assesment#5** | **Assesment#6** |
| **Mouse ID** | **CN264** | **CN264** | **CN264** | **CN264** | **CN264** | **CN264** |
| Gender | Male | Male | Male | Male | Male | Male |
| Age at assesment | 3 months | 4 months | 5 months | 6 months | 7 months | 8 months |
| Tremor | 0 | 0 | 0 | 0 | 0 | 0 |
| Body position | 0 | 0 | 0 | 0 | 0 | 0 |
| Gait | 0 | 0 | 0 | 0 | 0 | 0 |
| Grooming | 0 | 1 | 1 | 1 | 1 | 0 |
| Limb Tone | 0 | 0 | 0 | 0 | 0 | 0 |
| Weight score | 0 | 0 | 0 | 0 | 0 | 0 |
| **Total Score** | 0 | 1 | 1 | 1 | 1 | 0 |
| **Untreated** | **Assesment#1** | **Assesment#2** | **Assesment#3** | **Assesment#4** | **Assesment#5** | **Assesment#6** |
| **Mouse ID** | **CN265** | **CN265** | **CN265** | **CN265** | **CN265** | **CN265** |
| Gender | Male | Male | Male | Male | Male | Male |
| Age at assesment | 3 months | 4 months | 5 months | 6 months | 7 months | 8 months |
| Tremor | 0 | 0 | 0 | 0 | 0 | 0 |
| Body position | 0 | 0 | 0 | 0 | 0 | 0 |
| Gait | 0 | 0 | 0 | 0 | 0 | 0 |
| Grooming | 0 | 1 | 1 | 1 | 1 | 1 |
| Limb Tone | 0 | 0 | 0 | 0 | 0 | 0 |
| Weight score | 0 | 0 | 0 | 0 | 1 | 0 |
| **Total Score** | 0 | 1 | 1 | 1 | 2 | 1 |
| **Untreated** | **Assesment#1** | **Assesment#2** | **Assesment#3** | **Assesment#4** | **Assesment#5** | **Assesment#6** |
| **Mouse ID** | **CN266** | **CN266** | **CN266** | **CN266** | **CN266** | **CN266** |
| Gender | Male | Male | Male | Male | Male | Male |
| Age at assesment | 3 months | 4 months | 5 months | 6 months | 7 months | 8 months |
| Tremor | 0 | 0 | 0 | 0 | 0 | 0 |
| Body position | 0 | 0 | 0 | 0 | 0 | 0 |
| Gait | 0 | 0 | 0 | 0 | 0 | 0 |
| Grooming | 0 | 1 | 1 | 1 | 1 | 1 |
| Limb Tone | 0 | 0 | 0 | 0 | 0 | 0 |
| Weight score | 0 | 0 | 0 | 0 | 0 | 1 |
| **Total Score** | 0 | 1 | 1 | 1 | 1 | 2 |
| **Untreated** | **Assesment#1** | **Assesment#2** | **Assesment#3** | **Assesment#4** | **Assesment#5** | **Assesment#6** |
| **Mouse ID** | **CN267** | **CN267** | **CN267** | **CN267** | **CN267** | **CN267** |
| Gender | Male | Male | Male | Male | Male | Male |
| Age at assesment | 3 months | 4 months | 5 months | 6 months | 7 months | 8 months |
| Tremor | 0 | 0 | 0 | 0 | 0 | 0 |
| Body position | 0 | 0 | 0 | 0 | 0 | 0 |
| Gait | 0 | 0 | 0 | 0 | 0 | 0 |
| Grooming | 0 | 1 | 1 | 1 | 1 | 1 |
| Limb Tone | 1 | 0 | 0 | 0 | 0 | 0 |
| Weight score | 1 | 0 | 0 | 0 | 0 | 0 |
| **Total Score** | 2 | 1 | 1 | 1 | 1 | 1 |
| **Untreated** | **Assesment#1** | **Assesment#2** | **Assesment#3** | **Assesment#4** | **Assesment#5** | **Assesment#6** |
| **Mouse ID** | **CN268** | **CN268** | **CN268** | **CN268** | **CN268** | **CN268** |
| Gender | Male | Male | Male | Male | Male | Male |
| Age at assesment | 3 months | 4 months | 5 months | 6 months | 7 months | 8 months |
| Tremor | 0 | 0 | 0 | 0 | 0 | 0 |
| Body position | 0 | 0 | 0 | 0 | 0 | 0 |
| Gait | 0 | 0 | 0 | 0 | 0 | 0 |
| Grooming | 0 | 1 | 1 | 1 | 1 | 1 |
| Limb Tone | 0 | 1 | 0 | 0 | 0 | 0 |
| Weight score | 0 | 1 | 0 | 0 | 0 | 0 |
| **Total Score** | 0 | 3 | 1 | 1 | 1 | 1 |
| **Untreated** | **Assesment#1** | **Assesment#2** | **Assesment#3** | **Assesment#4** | **Assesment#5** | **Assesment#6** |
| **Mouse ID** | **CN269** | **CN269** | **CN269** | **CN269** | **CN269** | **CN269** |
| Gender | Male | Male | Male | Male | Male | Male |
| Age at assesment | 3 months | 4 months | 5 months | 6 months | 7 months | 8 months |
| Tremor | 0 | 0 | 0 | 0 | 0 | 0 |
| Body position | 0 | 0 | 0 | 0 | 0 | 0 |
| Gait | 0 | 0 | 0 | 0 | 0 | 0 |
| Grooming | 0 | 1 | 1 | 1 | 1 | 1 |
| Limb Tone | 0 | 0 | 0 | 0 | 0 | 0 |
| Weight score | 0 | 0 | 0 | 0 | 0 | 0 |
| **Total Score** | 0 | 1 | 1 | 1 | 1 | 1 |

## References

1 Chessum, N., Jones, K., Pasqua, E. & Tucker, M. Recent advances in cancer therapeutics. Progress in medicinal chemistry 54, 1–63, doi:10.1016/bs.pmch.2014.11.002 (2015).

2 Giannini, G., Cabri, W., Fattorusso, C. & Rodriquez, M. Histone deacetylase inhibitors in the treatment of cancer: overview and perspectives. Future Medicinal Chemistry 4, 1439–1460, doi:10.4155/Fmc.12.80 (2012).

3 Mottamal, M., Zheng, S. L., Huang, T. L. & Wang, G. D. Histone Deacetylase Inhibitors in Clinical Studies as Templates for New Anticancer Agents. Molecules 20, 3898–3941, doi:10.3390/molecules20033898 (2015).

4 Falkenberg, K. J. & Johnstone, R. W. Histone deacetylases and their inhibitors in cancer, neurological diseases and immune disorders (vol 13, pg 673, 2014). Nature Reviews Drug Discovery 14, 219–219, doi:10.1038/nrd4579 (2014).

5 Dokmanovic, M., Clarke, C. & Marks, P. A. Histone deacetylase inhibitors: Overview and perspectives. Molecular Cancer Research 5, 981–989, doi:10.1158/1541-7786.Mcr-07-0324 (2007).

6 Xu, W. S., Parmigiani, R. B. & Marks, P. A. Histone deacetylase inhibitors: molecular mechanisms of action. Oncogene 26, 5541–5552, doi:10.1038/sj.onc.1210620 (2007).

7 Di, X.-J., Han, D.-Y., Wang, Y.-J., Chance, M. R. & Mu, T.-W. SAHA Enhances Proteostasis of Epilepsy-Associated α1(A322D)β2γ2 GABA(a) Receptors. Chemistry & biology 20, 10.1016/j.chembiol.2013.1009.1020, doi:10.1016/j.chembiol.2013.09.020 (2013).

8 Yang, C. et al. Histone deacetylase inhibitors increase glucocerebrosidase activity in Gaucher disease by modulation of molecular chaperones. Proceedings of the National Academy of Sciences of the United States of America 110, 966–971, doi:10.1073/pnas.1221046110 (2013).

9 Calamini, B. & Morimoto, R. I. Protein Homeostasis as a Therapeutic Target for Diseases of Protein Conformation. Current Topics in Medicinal Chemistry 12, 2623–2640 (2012).

10 Alam, M. S., Getz, M. & Haldar, K. Chronic administration of an HDAC inhibitor treats both neurological and systemic Niemann-Pick type C disease in a mouse model. Science translational medicine 8, 326ra323, doi:10.1126/scitranslmed.aad9407 (2016).

11 Vanier, M. T. Niemann-Pick disease type C. Orphanet J Rare Dis 5, 16, doi:10.1186/1750-1172-5-16 (2010).

12 Carstea, E. D. et al. Niemann-Pick C1 disease gene: homology to mediators of cholesterol homeostasis. Science 277, 228–231 (1997).

13 Naureckiene, S. et al. Identification of HE1 as the second gene of Niemann-Pick C disease. Science 290, 2298–2301 (2000).

14 X. Xie, M. S. B., J. M. Shelton, J. A. Richardson, J. L. Goldstein, G. Liang. Amino acid substitution in NPC1 that abolishes cholesterol binding reproduces phenotype of complete NPC1 deficiency in mice. Proc. Natl. Acad. Sci. U.S.A. 108, 15330–15335 (2011).

15 Yu, T. & Lieberman, A. P. Npc1 acting in neurons and glia is essential for the formation and maintenance of CNS myelin. PLoS Genet 9, e1003462, doi:10.1371/journal.pgen.1003462 (2013).

16 Kennedy, B. E., Hundert, A. S., Goguen, D., Weaver, I. C. & Karten, B. Presymptomatic Alterations in Amino Acid Metabolism and DNA Methylation in the Cerebellum of a Murine Model of Niemann-Pick Type C Disease. The American journal of pathology 186, 1582–1597, doi:10.1016/j.ajpath.2016.02.012 (2016).

17 Alobaidy, H. Recent advances in the diagnosis and treatment of niemann-pick disease type C in children: a guide to early diagnosis for the general pediatrician. Int J Pediatr 2015, 816593, doi:10.1155/2015/816593 (2015).

18 Guillemot, N., Troadec, C., de Villemeur, T. B., Clement, A. & Fauroux, B. Lung disease in Niemann-Pick disease. Pediatric pulmonology 42, 1207–1214, doi:10.1002/ppul.20725 (2007).

19 Griese, M. et al. Respiratory disease in Niemann-Pick type C2 is caused by pulmonary alveolar proteinosis. Clinical genetics 77, 119–130, doi:10.1111/j.1399-0004.2009.01325.x (2010).

20 Lyseng-Williamson, K. A. Miglustat: a review of its use in Niemann-Pick disease type C. Drugs 74, 61–74, doi:10.1007/s40265-013-0164-6 (2014).

21 Nagral, A. Gaucher disease. Journal of clinical and experimental hepatology 4, 37–50, doi:10.1016/j.jceh.2014.02.005 (2014).

22 J. E. Wraith, D. V. E. Jacklin, L. Abel, H. Chadha-Boreham, C. Luzy, R. Giorgino, M. C. Patterson. Miglustat in adult and juvenile patients with Niemann-Pick disease type C: Long-term data from a clinical trial. Mol. Genet. Metab. 99, 351–357 (2010).

23 Pineda, M. et al. Clinical experience with miglustat therapy in pediatric patients with Niemann-Pick disease type C: a case series. Molecular genetics and metabolism 99, 358–366, doi:10.1016/j.ymgme.2009.11.007 (2010).

24 Liu, B. et al. Reversal of defective lysosomal transport in NPC disease ameliorates liver dysfunction and neurodegeneration in the npc1-/-mouse. Proc Natl Acad Sci U S A 106, 2377–2382, doi:10.1073/pnas.0810895106 (2009).

25 Ory, D. S. et al. Intrathecal 2-hydroxypropyl-beta-cyclodextrin decreases neurological disease progression in Niemann-Pick disease, type C1: a non-randomised, open-label, phase 1-2 trial. Lancet, doi:10.1016/s0140-6736(17)31465-4 (2017).

26 C. C. Pontikis, C. D. D., S. U. Walkley, F. M. Platt, D. J. Begley. Cyclodextrin alleviates neuronal storage of cholesterol in Niemann-Pick C disease without evidence of detectable blood–brain barrier permeability. J. Inherit. Metab. Dis. 36, 491–498 (2013).

27 Aqul, A. et al. Unesterified cholesterol accumulation in late endosomes/lysosomes causes neurodegeneration and is prevented by driving cholesterol export from this compartment. J Neurosci 31, 9404–9413, doi:10.1523/jneurosci.1317-11.2011 (2011).

28 Vite, C. H. et al. Intracisternal cyclodextrin prevents cerebellar dysfunction and Purkinje cell death in feline Niemann-Pick type C1 disease. Science translational medicine 7, 276ra226, doi:10.1126/scitranslmed.3010101 (2015).

29 Ramirez, C. M. et al. Weekly cyclodextrin administration normalizes cholesterol metabolism in nearly every organ of the Niemann-Pick type C1 mouse and markedly prolongs life. Pediatr Res 68, 309–315, doi:10.1203/00006450-201011001-0060410.1203/PDR.0b013e3181ee4dd2 (2010).

30 Ramirez, C. M. et al. Quantitative role of LAL, NPC2, and NPC1 in lysosomal cholesterol processing defined by genetic and pharmacological manipulations. Journal of lipid research 52, 688–698, doi:10.1194/jlr.M013789 (2011).

31 Kirkegaard, T. et al. Heat shock protein-based therapy as a potential candidate for treating the sphingolipidoses. Science translational medicine 8, 355ra118, doi:10.1126/scitranslmed.aad9823 (2016).

32 Maue, R. A. et al. A novel mouse model of Niemann-Pick type C disease carrying a D1005G-Npc1 mutation comparable to commonly observed human mutations. Human Molecular Genetics 21, 730–750, doi:10.1093/hmg/ddr505 (2012).

33 Fischer, A. H., Jacobson, K. A., Rose, J. & Zeller, R. Hematoxylin and eosin staining of tissue and cell sections. CSH protocols 2008, pdb.prot4986, doi:10.1101/pdb.prot4986 (2008).

34 Norwood, J., Franklin, J. M., Sharma, D. & D'Mello, S. R. Histone deacetylase 3 is necessary for proper brain development. J Biol Chem 289, 34569–34582, doi:10.1074/jbc.M114.576397 (2014).

35 Montgomery, R. L., Hsieh, J., Barbosa, A. C., Richardson, J. A. & Olson, E. N. Histone deacetylases 1 and 2 control the progression of neural precursors to neurons during brain development. Proc Natl Acad Sci U S A 106, 7876–7881, doi:10.1073/pnas.0902750106 (2009).

36 Volmar, C.-H. & Wahlestedt, C. Histone deacetylases (HDACs) and brain function. Neuroepigenetics 1, 20–27, doi:https://doi.org/10.1016/j.nepig.2014.10.002 (2015).

37 Venkatraman, A. et al. The histone deacetylase HDAC3 is essential for Purkinje cell function, potentially complicating the use of HDAC inhibitors in SCA1. Human Molecular Genetics 23, 3733–3745, doi:10.1093/hmg/ddu081 (2014).

38 Chitnis, T. & Weiner, H. L. CNS inflammation and neurodegeneration. The Journal of Clinical Investigation 127, doi:10.1172/JCI90609 (2017).

39 Ransohoff, R. M. How neuroinflammation contributes to neurodegeneration. Science 353, 777–783, doi:10.1126/science.aag2590 (2016).

40 Yanjanin, N. M. et al. Linear clinical progression, independent of age of onset, in Niemann-Pick disease, type C. American journal of medical genetics. Part B, Neuropsychiatric genetics : the official publication of the International Society of Psychiatric Genetics 153b, 132–140, doi:10.1002/ajmg.b.30969 (2010).

41 Alam, M. S. et al. Plasma signature of neurological disease in the monogenetic disorder Niemann-Pick Type C. J Biol Chem 289, 8051–8066, doi:10.1074/jbc.M113.526392 (2014).

42 Alam, M. S. et al. Genomic Expression Analyses Reveal Lysosomal, Innate Immunity Proteins, as Disease Correlates in Murine Models of a Lysosomal Storage Disorder. Plos One 7, doi:ARTN e4827310.1371/journal.pone.0048273 (2012).

43 Porter, F. D. et al. Cholesterol oxidation products are sensitive and specific blood-based biomarkers for Niemann-Pick C1 disease. Science translational medicine 2, 56ra81, doi:10.1126/scitranslmed.3001417 (2010).

44 Jiang, X. et al. Development of a bile acid-based newborn screen for Niemann-Pick disease type C. Science translational medicine 8, 337ra363, doi:10.1126/scitranslmed.aaf2326 (2016).

45 Liu, Y. & Bankaitis, V. A. Phosphoinositide phosphatases in cell biology and disease. Prog Lipid Res 49, 201–217 (2010).

46 Subramanian, K., Rauniyar, N., Lavallee-Adam, M., Yates, J. R., 3rd & Balch, W. E. Quantitative Analysis of the Proteome Response to the Histone Deacetylase Inhibitor (HDACi) Vorinostat in Niemann-Pick Type C1 disease. Molecular & cellular proteomics : MCP, doi:10.1074/mcp.M116.064949 (2017).

